# H2AFX and MDC1 Protect Genomic Integrity in Male Germ Cells by Promoting Recombination and Activation of the Recombination-Dependent Checkpoint

**DOI:** 10.1101/235085

**Authors:** Erika Testa, Daniela Nardozi, Cristina Antinozzi, Monica Faieta, Stefano Di Cecca, Cinzia Caggiano, Tomoyuki Fukuda, Elena Bonanno, Lou Zenkun, Andros Maldonado, Ignasi Roig, Monica Di Giacomo, Marco Barchi

**Author notes:** Equally contributing authors (Cristina Antinozzi, Monica Faieta). Current address: Department of Movement, Human and Health Sciences, University of Rome Foro Italico, 00135 Rome, Italy (Cristina Antinozzi). Corresponding author: Marco Barchi, Ph.D., Department of Biomedicine and Prevention, University of Rome Tor, Vergata, Via Montpellier n.1, 00133, Rome, Italy.

## Abstract

In somatic cells, *H2afx* and *Mdc1* are close functional partners in DNA repair and damage response. However, it is not known whether they are also involved in the maintenance of genome integrity in meiosis. By analyzing chromosome dynamics in *H2afx^-/-^* spermatocytes, we found that synapsis of the autosomes and X-Y chromosomes were impaired in a relevant fraction of cells. Such defect correlated with an abnormal recombination profile. Conversely, *Mdc1* was dispensable for the synapsis of the autosomes, and only played a minor role in X-Y synapsis, relatively to *H2afx*. This suggested that those genes have non-overlapping functions in chromosome synapsis. However, we observed that both genes play a similar role in the assembly of MLH3 onto chromosomes, a key step in crossover formation. Moreover, we showed that *H2afx* and *Mdc1* cooperate in promoting the activation of the recombination-dependent checkpoint, a mechanism that restrains the differentiation of cells with unrepaired DSBs. This occurs by a mechanism that involves P53. Overall, our data showed that, in male germ cells, *H2afx* and *Mdc1* promote the maintenance of genome integrity.

## Introduction

The production of gametes with an intact haploid genome is critical for the prevention of birth defects. It depends on the formation of crossovers (COs), the chromosome connections that ensure faithful segregation during metaphase I (Keeney, 2008). CO formation is achieved by meiotic recombination, a process that is initiated with the introduction of double-strand breaks (DSBs) by the SPO11 protein. DSBs are potentially dangerous, because they introduce opportunities for generating gross chromosomal rearrangements that may be detrimental to germ cell genome integrity (Kim et al., 2016). In addition, as in many organisms, in mammals SPO11-induced DNA damage is coupled with synapsis of homologous chromosomes (homologs) (Bolcun-Filas and Schimenti, 2012), failure to recombine can cause aneuploidy and lead to the development of an embryo with an unbalanced chromosome number. In order to avoid such dangerous outcomes, surveillance mechanisms (or “checkpoints”) sense defects in meiotic recombination/synapsis, and arrest the progression of cells with unresolved problems.

In male mice, the meiotic progression of germ cells is monitored by two main distinct mechanisms: the “sex body-deficient checkpoint” and the “recombination-dependent checkpoint”. The former is triggered in synapsis-defective cells by faulty transcriptional silencing of X-Y associated genes that are toxic to male germ cells (Mahadevaiah et al., 2008, Burgoyne et al., 2009, Royo et al., 2010). The latter is instead activated in cells that have an abnormal recombination profile. In pachynema cells, it co-ordinates DSBs repair with meiotic progression, thus causing a delay of differentiation until a sufficient number of DSBs have been repaired (e.g. as in *Tex19.1^-/-^* spermatocytes (James H Crichton, 2017)) or, ultimately, apoptosis if DSBs cannot be properly resolved (e.g. if they lack TRIP13, an AAA+ ATPase required to timely complete meiotic recombination (Pacheco et al., 2015)). Genetic evidence indicates that requirements of the sex body-deficient checkpoint and the recombination-dependent checkpoint are different. Activation of the former relies on many different genes (Turner, 2015), including *H2afx* (nucleosomal variant histone *H2afx*) and *Mdc1* (mediator of DNA damage checkpoint protein 1). Silencing of X-Y chromosomes occurs following phosphorylation of H2AFX on Ser 139 (which is referred to as γH2AFX) by the ataxia telangiectasia and Rad3-related protein (ATR) (Fernandez-Capetillo et al., 2003, Mahadevaiah et al., 2008), and spreading of such signal over the sex body domain through MDC1 (Lou et al., 2003, Stucki et al., 2005, Ichijima et al., 2011). Instead, activation of the recombination-dependent checkpoint depends on the activity of the Mre11 complex, the DNA damage sensor kinase ATM (ataxia-telangiectasia mutated), its downstream target CHK2 (Pacheco et al., 2015) and P53/TAp63 family members (Marcet-Ortega et al., 2017).

In somatic cells, checkpoint response to DNA damage entails a functional interplay between H2AFX, MDC1 and ATM. ATM-mediated formation of γH2AFX is one of the earliest responses to damage (Rogakou et al., 1998, Rogakou et al., 1999). MDC1 binding onto γH2AFX promotes ATM accumulation at the sites of DNA damage. This amplifies the DNA damage signals (Lou et al.,2006), including activation of the ATM-target P53 (Canman et al., 1998, Khanna et al., 1998, Banin et al., 1998). Another key response to DNA damage response is DNA repair. In this respect, γH2AFX and MDC1 have been found to promote DNA repair by homologous recombination, (Unal et al., 2004, Xie et al., 2004, Sonoda et al., 2007, Zhang et al., 2005, Xie et al., 2007) a process that ensures repair of DNA damage and synapsis of the homologs in mammalian meiosis. However, whether H2AFX and MDC1 play a function in checkpoint response and meiotic recombination in meiotic cells is unclear. Genetic evidence indicates that, in *Mdc1* mutant spermatocytes, synapsis of the autosomes progresses normally, thus suggesting no overt function in recombination (Ichijima et al., 2011). Whether H2AFX is needed for proper synapsis of the autosomes remains unclear. In fact, while according to some authors the synapsis of the autosomes failed (Moens et al., 2007), others reported no defect (Celeste et al., 2002, Fernandez-Capetillo et al., 2003, Mahadevaiah et al., 2008). Interestingly, in spermatocytes of both *Mdc1^-/-^* and *H2afx^-/-^*, X-Y synapsis fails with some degree (Fernandez-Capetillo et al., 2003, Ichijima et al., 2011). This indicates that, somehow, those genes support recombination between sex chromosomes. However, the mechanisms underlying this defect have not been understood.

By comparing the phenotype of *H2afx* and *Mdc1* mutants, we demonstrated that DSBs form normally in both mutants. However, in absence of *H2afx*, formation of MSH4 foci and synapsis of the autosomes is impaired. On the contrary, such defects are not observed in *Mdc1^-/-^* cells, which indicates that *H2afx* and *Mdc1* are functionally separated in recombination-mediated synapsis. Accordingly, X-Y asynapsis is much higher in *H2afx^-/-^* spermatocytes than in *Mdc1* mutant cells. Therefore, *Mdc1* performs a minor function in recombination-mediated synapsis of the sex chromosomes. Beside such differences, H2AFX and MDC1 play a similar role in promoting the assembly of MLH3 foci (a marker of COs formation). This points to a role of said genes in the pathway that enables stable interaction and segregation of the homologs at metaphase I. Furthermore, in line with the function of H2AFX and MDC1 in controlling cell cycle in somatic cells with damaged DNA, we showed evidence that both H2AFX and MDC1 also play a role in the activation of the recombination-dependent checkpoint, promoting the activation of P53.

Overall, these data highlight that *H2afx* and *Mdc1* sustain genome integrity preservation of male germ cells, by both promoting proper processing of DSBs, and delaying the progression of cells with unrepaired damage.

## Results

### *H2afx* and *Mdc1* are dispensable to regulate formation of nucleus-wide DSBs

In mammalian meiocytes, the activity of the PI3K-like kinase ATM is essential to constrain the formation of nucleus-wide DSBs by SPO11 (Lange et al., 2011). Therefore, targets of ATM are expected to exert control of DSBs formation. In mouse meiosis, H2AFX is phosphorylated by ATM at leptonema and zygonema, as a response to DSBs formation (Barchi et al., 2005, Bellani et al., 2005). This suggests that H2AFX might be part of the regulatory mechanism that control the function of SPO11. In addition, the γH2AFX binding partner MDC1 is also a target of ATM (Matsuoka et al., 2007, Lavin and Kozlov, 2007, Jungmichel et al., 2012). However, whether H2AFX or MDC1 regulates the number of DSBs in meiosis is unknown.

To assess nucleus-wide DSBs formation in *H2afx^-/-^* males, we quantified SPO11-oligonucleotide complexes that are a byproduct of meiotic DSBs formation (Pan et al., 2011, Neale et al., 2005, Lange et al., 2011). In mice, the first wave of spermatogenesis is semi-synchronous and begins around six days post-partum (dpp). Pachytene stage cells appear approximately at 12-14 dpp, followed by the diplotene stage at 18 dpp. *H2afx^-/-^* spermatocytes undergo apoptosis by mid-pachynema (Fernandez-Capetillo et al., 2003, Mahadevaiah et al., 2008). Therefore, in our experiment, in order to minimize changes in the cellular composition of the testis relative to wild-type mice, we used 12.5 dpp juvenile mice. Quantification of SPO11-oligonucleotide complexes steady-state level in a single experiment revealed a small weakening of the signal in *H2afx^-/-^* testes compared to the control (71,2% of wild type) (Fig. 1A). This may reflect stochastic variability in the samples, or alternatively be due to a reduction of DSBs in the mutant. In order to make a distinction between those two options, we sought to determine the number of DSBs by quantifying DMC1 foci in leptonema and early/mid zygonema cells. DMC1 is a meiosis-specific member of the bacterial RecA protein family, which stabilizes strand exchange intermediates to promote DSBs repair (Bishop et al., 1992, Yoshida et al., 1998). Cytologically, it can be seen as foci along meiotic chromosome axes, and has been broadly used as a surrogate marker for DSBs formation (Lam and Keeney, 2015). We detected no differences between the genotypes, which suggests that in absence of *H2afx*, SPO11 activity is not altered (Fig.1 B and S1 table). Similarly, DMC1 foci number in *Mdc1* knockout spermatocytes were not different compared to controls (Fig. 1B). Hence, we concluded that albeit ATM plays in meiosis a key role in the homeostatic control of global DSB numbers (Lange et al., 2011), such function does not rely on H2AFX or MDC1.

**Fig.1.**
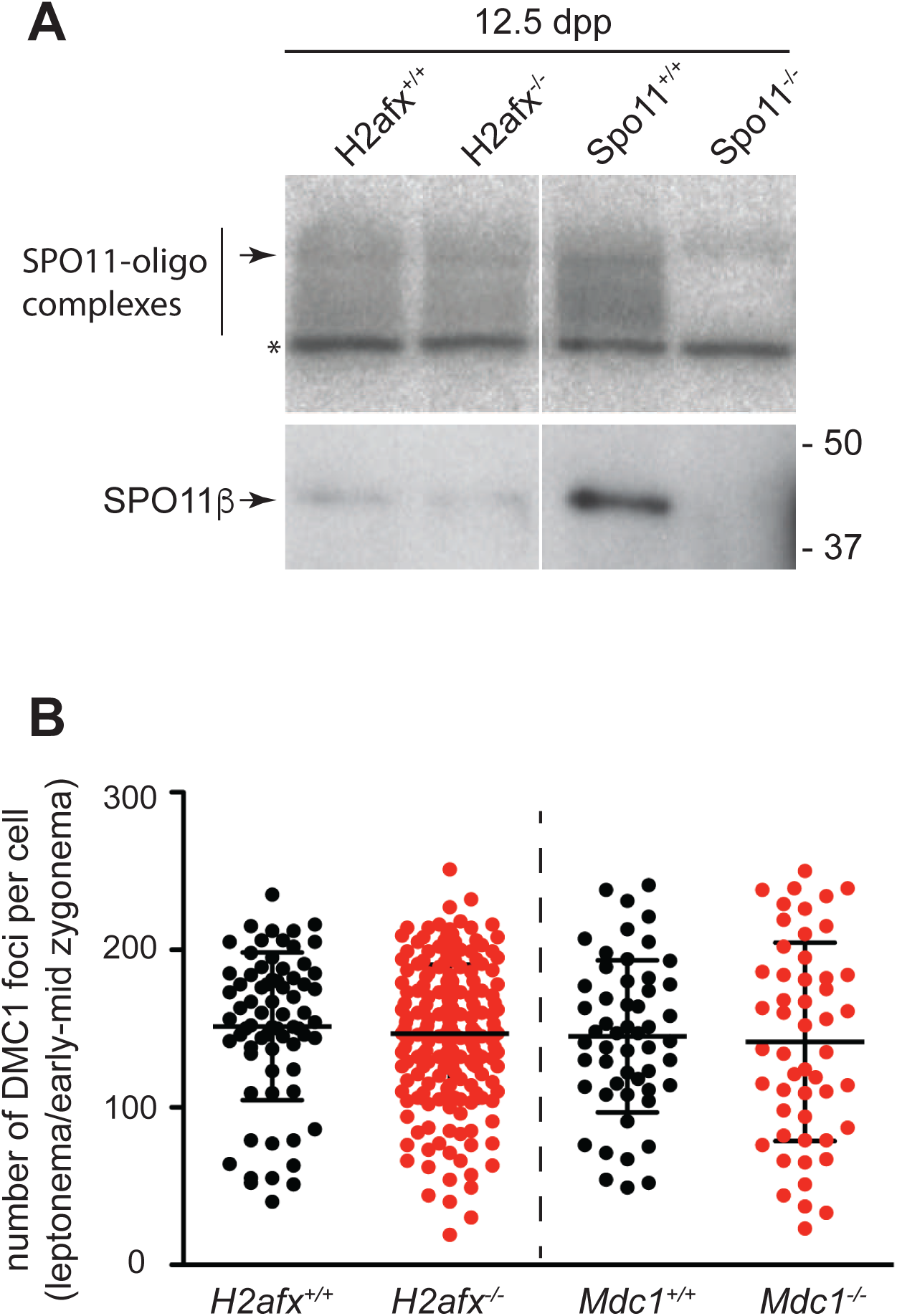
*H2Afx* and *Mdc1* are dispensable for the formation and homeostatic control of DSBs. (A) Autoradiography of SPO11-oligonucleotide complexes (upper panel) and anti-Spo11 western detection (lower panel) isolated by immunoprecipitation of juvenile mice testis. One mouse (two testis) per genotype was analyzed. Asterisk= non-specific terminal transferase labeling; arrowheads represent migration position of immunoglobulin heavy chain. Molecular mass is given in kilodalton. (B) Quantification of DMC1 foci number of the indicated genotypes of 12.5 dpp juvenile mice. Three to four mice were analyzed for each genotype. Each *dot* in the graph indicates the number of DMC1 foci per nucleus; the error bars are means and standard deviation (for more details, see S1 table).

### Defective X-Y chromosome synapsis in *H2afx^-/-^* and *Mdc1^-/-^* cells do not rely on reduced formation of DSBs in the PAR

During meiotic prophase I, synapsis of the homologs occurs gradually and it is completed by pachynema. Cytologically, synapsis can be monitored by following the deposition of proteins of the lateral (e.g. SYCP3) and central (e.g. SYCP1) elements of the Synaptonemal Complex (SC), by immunofluorescence on spread chromosomes. In wild-type mice at pachynema, the autosomes are fully synapsed along their lengths; conversely, X-Y chromosomes synapsis is restricted to the pseudo autosomal region (PAR), a small region of homology between them (Perry et al., 2001). As the PAR is small, the X-Y chromosome pair is very sensitive to perturbations of the synaptic process. Confirming a previous report (Fernandez-Capetillo et al., 2003), we found that a lack of *H2afx* causes X-Y chromosomes synapsis defect in about 80% of the pachynema cells in juvenile males (Fig. 2A and S1A Fig.). This indicates that DSBs formation and/or recombination within the PAR are perturbed. However, the mechanisms underlying this defect are not known.

**Fig.2.**
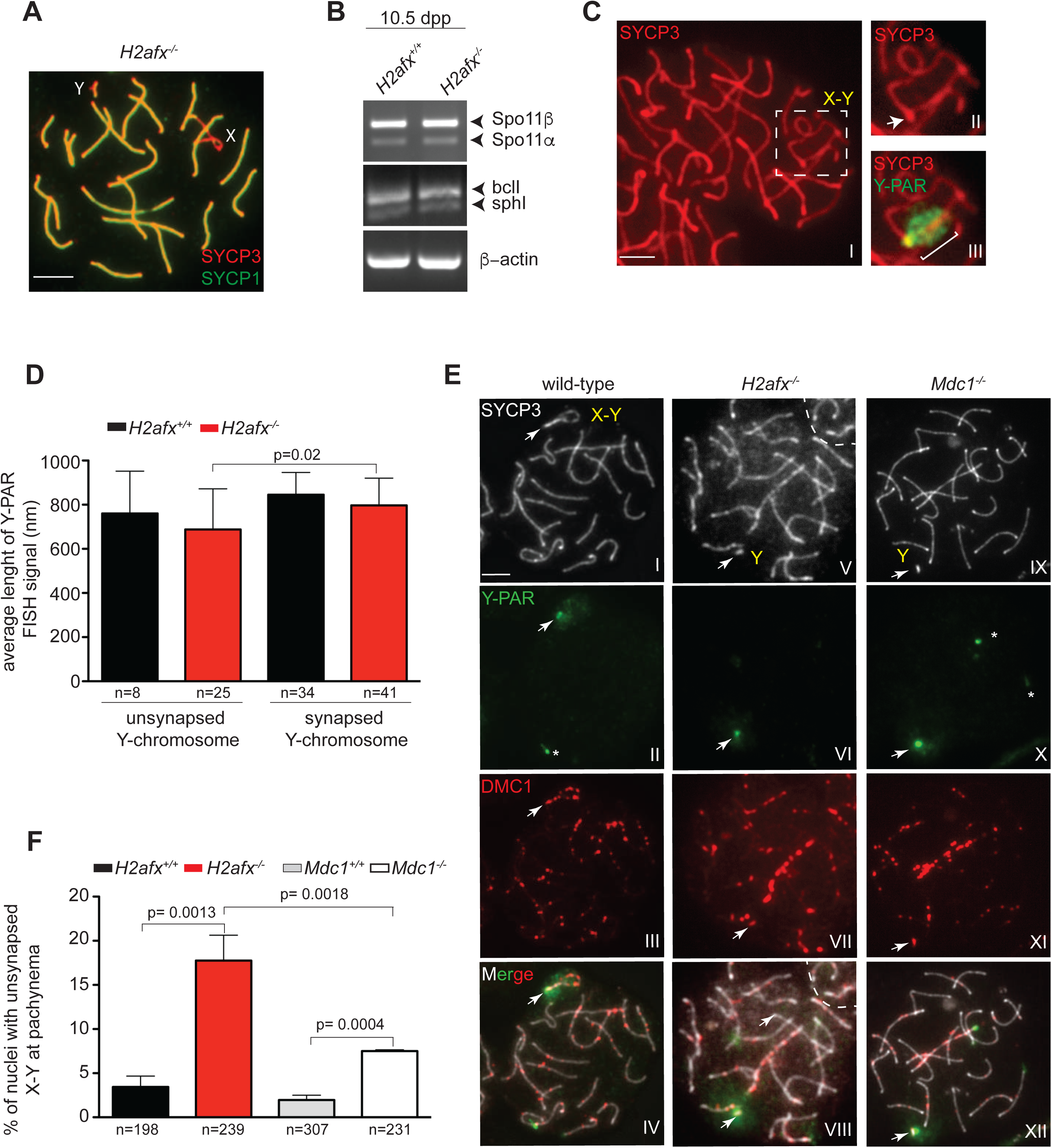
X-Y chromosome asynapsis in *H2afx^-/-^* and *Mdc1^-/-^* cells does not rely on reduced formation of DSBs in the PAR. (A) Representative nuclear spread in the pachytene stage of 12.5 dpp *H2afx^-/-^* spermatocyte labeled with anti-SYPC3 and anti-SYCP1. Scale bar represents 5 μm. (B) Representative RT-PCR analysis of the expression of *Spo11* splicing variant in the indicated genotypes of juvenile mice. Level of β-actin was used as loading control. (C) Representative image of synapsed X-Y chromosomes in wild-type spermatocytes at early-pachytene stage (panel I). SYCP3 (red) marks the axial element of the synaptonemal complex. X-Y chromosomes are encircled by the dotted square. Magnification in panel II shows the X-Y chromosomes. The arrow points to the PAR region. Magnification in panel III shows the FISH signal (green) identifying the PAR region. The white bar indicates the extension of the PAR-FISH signal that is measured as aproxy for chromatin loop length measure. The magnification bar in I represents 5μM. (D) Quantification of the average length of Y-PAR FISH signal among spermatocytes of indicated genotypes, with unsynapsed and synapsed Y-chromosome (*H2afx* ^+/+^ n= 3 mice; *H2afx ^-/-^* n= 3 mice). Error bars = mean±SD; p= p value. (E) Representative early pachytene spermatocytes of indicated genotypes, labeled with anti-SYCP3 (I,V, IX), Y-PAR probe (II,VI, X) and anti-DMC1 antibody (III,VII,XI). White arrows indicate Y-chromosome, while asterisks indicate a non-specific signal of the FISH. Merge images (IV, VIII, XII) show the localization of DMC1 foci within the Y-PAR region. Three adult mice were tested for each genotype. (F) Quantification of nuclei with unsynapsed X-Y chromosomes at the pachynema stage in the indicated genotypes. Three adult mice were tested for each genotype. Error bars = mean±SD; p= p value.

In mice, DSB formation within the PAR requires proper expression of *Spo11* splicing variants (Kauppi et al., 2011). To investigate proper *Spo11* splice variants expression in *H2afx* mutant, we amplified the cDNA of *H2afx^-/-^* and wild-type testis, using *Spo11* isoform-specific primers. For this experiment, we used 10.5 dpp juvenile mice because at this age the cellular composition of the testis is expected to be similar (Fernandez-Capetillo et al., 2003, Mahadevaiah et al., 2008). We observed that *Spo11β, Spo11α, bcll* and *sphI* isoforms (Keeney et al., 1999) were all expressed in the mutant, at a level comparable with that of control (Fig. 2B). This indicates that the X-Y chromosome asynapsis in the mutant is unrelated to changes in *Spo11* splicing pattern.

In addition to the proper expression of *Spo11* splicing variants, DSBs formation within the PAR relies on the structural characteristics of the PAR. PAR axes in mice are disproportionally long relative to DNA length. This results in shorter chromatin loops, with consequent increase in DSB potentials (Kauppi et al., 2011). H2AFX is a structural component of chromatin. We therefore reasoned that its absence might cause an increase in the size of PAR-associated chromatin loops. To evaluate this possibility, by looking at chromosome spread preparations, we compared the PAR chromatin loop size of wild-type and *H2afx^-/-^* spermatocytes in early pachynema nuclei. With a view to identifying early substages of pachynema, we co-stained the chromosomes with an anti-DMC1 antibody, a marker that, in normal cells, persists at relatively high levels up to early pachynema. As a proxy for loop size, we measured fluorescence in situ hybridization (FISH) signal extension from axes of cells, using a PAR-specific probe (Fig. 2C). Under our experimental conditions, we could always unambiguously distinguish the Y-PAR, while the X-PAR signals were often weak and sometimes not visible (S1 B). Therefore, we focused our analyses on the Y-PAR. We observed that the loop size of unsynapsed Y-chromosomes of *H2afx^-/-^* cells at early/midpachynema had not increased compared to that measured in synapsed Y-chromosomes of wild-type or *H2afx^-/-^* cells (Fig. 2D). Actually, we rather measured a slight decrease, indicating that the asynapsis of the X-Y chromosomes is not linked to changes in chromatin loop organization. If our observations are correct, we assume that in *H2afx* spermatocytes, DSB occurs within the Y-PAR, at the wild-type level. To test this hypothesis, using DMC1 as a marker, we analyzed DSB formation within the PAR. Our analyses revealed that in the mutant, a DMC1 focus was observed within the Y-PAR of asynapsed chromosomes at early pachynema in 66.6±9% (n=30) of the nuclei (Fig. 2E I-VIII). This percentage closely matches that of the nuclei with Y-PAR associated foci at late zygonema in wild-type (Kauppi et al., 2011), indicating that a lack of H2AFX does not affect DSB formation within the PAR. In the absence of any defect in DSB formation, X-Y chromosome synapsis failure suggests that processing of PAR-associated DSBs is defective.

Previous findings have shown that, in addition to *H2afx,* X-Y synapsis also requires *Mdc1* (Ichijima et al., 2011). To understand whether in *Mdc1^-/-^* cells X-Y asynapsis rely on a defect in DSB formation within the PAR, we quantified the number of PAR-associated DMC1 foci. Similarly to what was observed in *H2afx* null spermatocytes, DSBs were present within the Y-PAR in about 66±10% (n=28) of cases (Fig. 2E IX-XII), thus ruling out a defect in DSB formation. Interestingly, the side-by-side analyses of frequency of X-Y chromosome asynapsis in adult mice from both *H2afx* and *Mdc1* mutants revealed that, although in *Mdc1^-/-^* spermatocytes the percentage of asynapsed chromosomes increased significantly relatively to the wild-type, such an increase was much smaller if compared to that observed in *H2afx^-/-^* cells (Fig. 2F). This indicates that, although both genes play a role in promoting the synapsis of sex chromosomes, the *H2afx* function predominates over that of *Mdc1*.

Overall, the aforesaid observations demonstrate that *H2afx* and *Mdc1* are not necessary for the formation of DSBs within PAR. However, both genes play a role in promoting recombination in this region, probably through mechanisms that are only partially overlapping (see discussion).

### *H2afx^-/-^* spermatocytes display defects in homologous synapsis of the autosomes

The above observation that in *H2afx^-/-^* mutant recombination is likely to be defective in most X-Y chromosomes, suggests that global recombination-mediated synapsis of the autosomes might also be disrupted. To challenge this statement, we sought to study autosome dynamics during prophase I. In mice, the synapsis of the autosomes is initiated at zygonema. Cytologically, progression from leptonema to early zygonema is characterized by the assembly of SYCP1, that occurs while the elongation of the axial (now referred to as lateral) element containing SYCP3 has already begun (Fig. 3A I). As cells progress to mid-zygonema, the elongation of the chromosome axes is completed, and chromosome synapsis further extended coordinately in most of the chromosomes, up to 50% of chromosome length (Fig. 3A II). At late-zygonema, homologous synapsis extends over 50% of chromosome length (Fig. 3AIII), and it is completed at pachynema (Fig. 3A IV).

**Fig.3.**
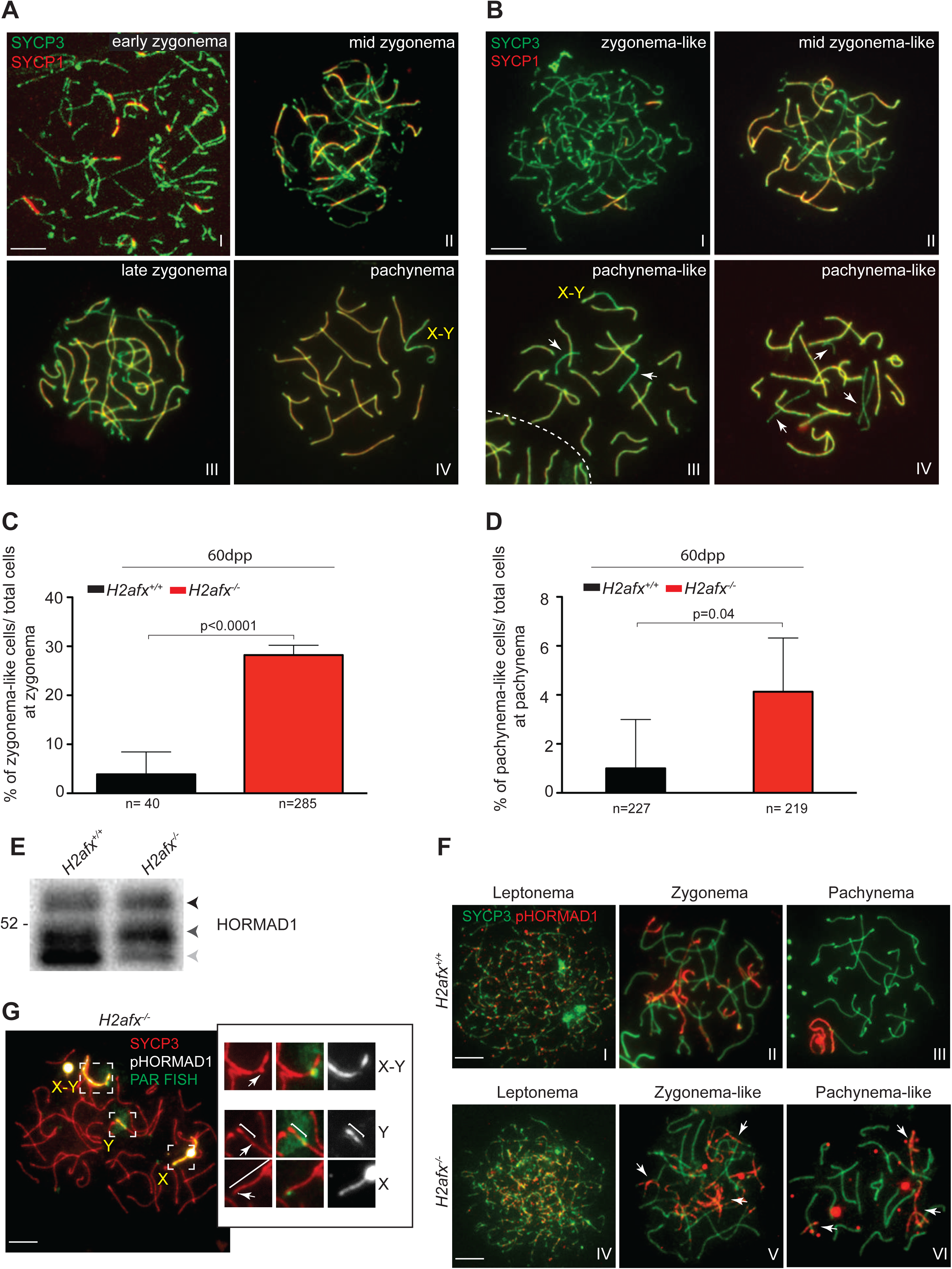
In *H2afx ^-/-^* spermatocytes homologous synapsis of the autosomes are defective. (A-B) Representative images of nuclear spreads of the indicated stages, from wild-type and *H2afx^-/-^* spermatocytes labeled with anti-SYCP3 and anti-SYCP1 antibodies. Arrows point axes of unsynapsed chromosomes. Scale bar is 5*μ*m. (C) Quantification *of* zygonema-like cells within zygonema class in 60 dpp mice of indicated genotypes (*H2afx*^+/+^ n= 3, *H2afx^-/-^ n*= 4 mice tested). (D) Quantification of pachynema-like cells within pachynema cells class in 60 dpp mice of indicated genotypes. (*H2afx* ^+/+^ *n*= 3; *H2afx ^-/-^ n*= 4 mice tested). Error bars are mean±SD; p= p value. (E) Westernblot analyses of testisinsoluble nuclear extracts from 60dpp mice testis of the indicated genotypes. Slow (phosphorylated) migrating forms of HORMAD1 are indicated by black and dark-grey arrows. One mouse per genotype was analyzed. Molecular mass is given in kilodaltons. (F) Representative images of nuclear spreads of spermatocytes from the indicated stages and genotypes, labeled with anti-SYCP3 and anti-pHORMAD1. Arrows point axes of unsynapsed homologs. (G) Representative image of *H2afx ^-/-^* spread spermatocytes labeled with anti-SYCP3 (red), anti-pHORMAD1 (white) and FISH (green) to identify the Y-PAR probe. Right panel, magnifications of dotted squares regions representing the pattern of pHORMAD1 in synapsed (upper panel) and unsynapsed sex chromosomes (lower panels). White bars indicate the extension of Y and X chromosomes. Scale bars are 5*μ*m.

A detailed analysis of autosome synapsis in *H2afx^-/-^* spermatocytes from adult mice revealed that a relevant fraction of cells at zygonema displayed defects in chromosome synapsis of the autosomes (henceforth referred to as zygonema-like cells) (Fig. 3 B-C), with different characteristics. Specifically, 23% of zygotene-stage cells displayed signs of delayed initiation of synapsis or asynchrony of synapsis among different homolog pairs. The former appeared as nuclei with fully elongated SYCP3-positive axes, with very limited synapsis (Fig. 3 B I). The latter, were instead cells characterized by the co-presence of multi fully-asynapsed bivalents in the presence of at least two fully-synapsed homolog pairs (Fig. 3B II). In addition to such global defects in the initiation or coordinate extension of chromosome synapsis, in a restricted population of spermatocytes at zygonema (5.3%), we found cells with interstitial asynapsis (a “bubble”) (S2 I Fig.) and cells displaying non-homologous synapsis (S2 II-III Figs). Non-homologous synapsis and “bubbles” were also observed concomitantly with the development of asynchronous synapsis in 5.4% of the cases.

There were also synaptic abnormalities at pachynema (Fig. 3D). Pachynema-like spermatocytes appeared as cells with only one or two pairs of unsynapsed autosomes in the presence of either synapsed (Fig. 3B III) or asynapsed sex chromosomes (Fig. 3B IV). The percentage of pachynema-like cells in 12.5 dpp juvenile mice was higher (9.6±6%, n=65) than that observed in the adults. This is plausibly due to the fact that, at that stage, most cells are still at early pachynema, a developmental stage that precedes their apoptotic elimination (Fernandez-Capetillo et al., 2003, Mahadevaiah et al., 2008, Royo et al., 2010).

The presence of asynapsed chromosomes at zygonema and pachynema in the mutant indicated that H2AFX might be required to initiate synapsis and promote chromosome pairing of the autosomes. However, an alternative interpretation is that H2AFX might be needed to prevent desynapsis caused by precocious dissociation of the homologs. To distinguish between these two options, we leveraged the different phosphorylation status of the HORMA (Hop1, Rev7 and Mad2) domain 1 (HORMAD1) protein on chromosome axes through prophase-I. HORMAD1 is a mammalian member of an evolutionarily preserved family of meiotic axis proteins, related to Hop1 in yeast (Wojtasz et al., 2009, Fukuda et al., 2010). In mice, it binds preferentially to chromosome axes where homologs are not synapsed, that is, axes prior to synapsis (un-synapsed) and axes where the SC has disassembled after the completion of synapsis (de-synapsed). However, protein phosphorylation at Ser^375^ (an ATM/ATR kinases consensus site) occurs onto un-synapsed, but not de-synapsed chromosomal regions (Fukuda et al., 2012). This makes HORMAD1 pSer^375^ (pHORMAD1) a reliable marker to distinguish between chromosomes showing asynapsis or de-synapsis defects. We first asked whether a lack of H2AFX impairs HORMAD proteins deposition. The analyses showed that HORMAD1, as well as HORMAD2, localize onto chromosomes with a pattern that is similar to that of control (S3 A-B Figs). To analyze whether HORMAD1 phosphorylation occurs normally in the absence of *H2afx*, we then performed a HORMAD1 immunobloting experiment of mouse testis insoluble nuclear extracts. As reported previously (Fukuda et al., 2012), this technique allows identifying the global phosphorylation status of chromosome-associated proteins. We observed that the phosphorylated forms of HORMAD1 (mid-grey and dark arrows head in Fig. 3E) were bound to chromosomes, thus indicating that H2AFX is dispensable for HORMAD1 phosphorylation. Similarly, this was observed for HORMAD2 and the other chromosome axis components SYCP2, STAG3, REC8 and SMC3 (see S3 C-D Figs). We then sought to specifically examine the chromosomal localization of HORMAD1 phosphorylation at Ser^375^ by using meiotic chromosome spread preparations. In the wild-type, pHORMAD1 staining followed the pattern described in literature (Figs 3F I-III) (Fukuda et al., 2012). In the *H2afx* mutant, asynapsed chromosomes of zygonema-like and pachynema-like cells were always phosphorylated (Figs 3F V-VI). From this observation we inferred that autosomes failed to initiate synapsis. To understand if such a defect also affects sex chromosomes, we analyzed pHORMAD1 staining pattern within the PAR. We observed that, in asynapsed X-Y chromosomes, the PAR was always phosphorylated (Fig. 3G). This indicates that, as for the autosomes, X-Y chromosomes fail to synapse.

It is worth highlighting that, as it has previously described by others (Ichijima et al., 2011), we observed that in *Mdc1^-/-^* spermatocytes, autosomes synapsis was substantially normal (cells with abnormal synapsis of the autosomes were: *Mdc1^-/-^*: 1±1% n=289; wt: 0% n=279), thus excluding any significant role for this gene in this process.

Overall, the aforementioned findings demonstrated that *H2afx* mutant cells suffered from a defect in the proper establishment of a physical interaction between autosomes.

### A fraction of *H2afx^-/-^* spermatocytes are eliminated by apoptosis before the completion of autosomal synapsis

In mice, spermatogenesis occurs within the seminiferous tubules in which, based on the specific germ cell association within each cross section, twelve epithelial stages (I-XII) can be identified (Russell, 1990, Ahmed and de Rooij, 2009). In the context of seminiferous epithelium, the lifespan of a spermatocyte that is unable to complete recombination and synapsis of the autosomes (e.g., if they lack the strand-ex-change protein DMC1 (Pittman et al., 1998, Yoshida et al., 1998)) is extended until the cells reach epithelial stage IV, a stage equivalent to mid-pachynema in wild-type (Barchi et al., 2005, Mahadevaiah et al., 2008, Burgoyne et al., 2009). Confirming previous observations (Mahadevaiah et al., 2008), our histological analyses of *H2afx^-/-^* mice testis revealed that, although a minority of cells escaped the checkpoint (De Rooij D.G., personal communication), most spermatocytes were eliminated by apoptosis at the epithelial stage IV (Fig. 4A). However, whether cells undergoing apoptosis reach stage IV and die before completion of autosome synapsis is currently unknown. To test this possibility, we performed a terminal deoxynucleotidyl transferase dUTP nick end labeling (TUNEL) on cells stained for SYCP3/SYCP1. As we observed in another experimental setting (Faieta et al., 2015), we noticed that strongly (most abundant) TUNEL-positive cells were too shrunken to be informative for the morphological analyses (S4 Fig.). This is likely because TUNEL marks late events of the apoptotic process. However, along with strongly TUNEL-positive cells, we identified a fraction of more weakly stained TUNEL-positive nuclei, where spermatocyte morphology was still preserved. The analyses of the latter sub-population revealed that, in addition to cells where autosomes were fully paired along their length (i.e. pachytene-stage spermatocytes), the TUNEL-positive population from *H2afx^-/-^* mice also contained a relevant fraction of cells where synapsis of the autosomes were not yet completed, i.e. zygotene and zygotene-like stage cells (Figs 4 B-C). Among them, the great majority were nuclei with an extension of synapsis that is typical of cells at early and mid-zygonema. This indicates that, in those nuclei, autosome synapsis was initiated but, just as it was observed in mutants with defects in recombination/synapsis of the autosomes (e.g. see (Barchi et al., 2005)), it could not be completed before reaching stage IV. To strengthen this observation, we also analyzed the morphology of TUNEL-positive nuclei in *Mdc1^-/-^* mutant spermatocytes. In agreement with the observation that a lack of *Mdc1* does not significantly impact autosomes synapsis (Ichijima et al., 2012), we observed that most cells in this mutant apoptose (at stage IV (Ahmed and de Rooij, 2009)) after completing autosomes synapsis (Fig. 4B). From these observations, we concluded that, in addition to defects in coordinate synapsis of the autosomes (see above), spermatocytes from *H2afx^-/-^* mice suffer from a defect in timely completion of autosome synapsis, due to a mechanism that does not involve *Mdc1*.

**Fig. 4.**
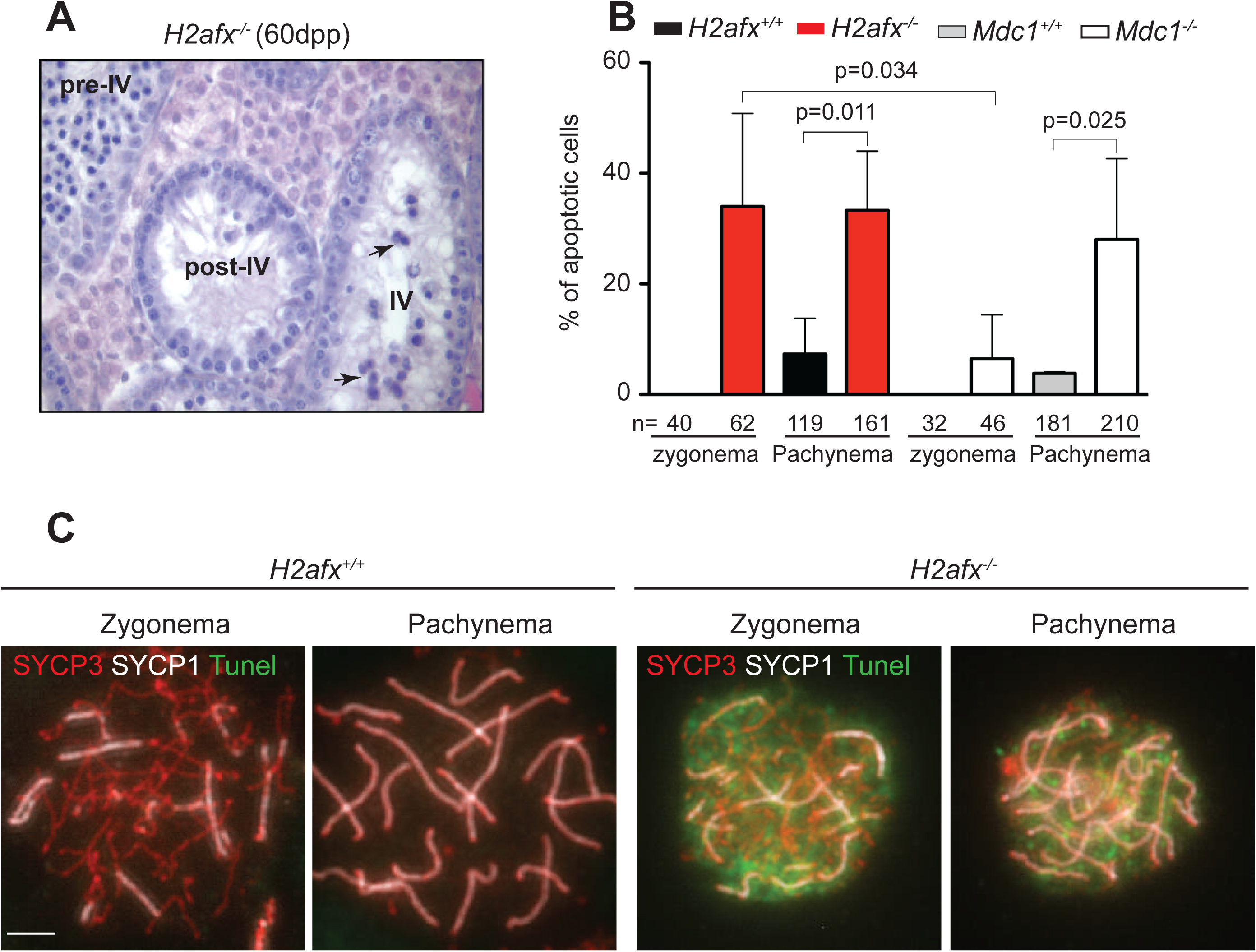
A fraction of *H2afx ^-/-^* spermatocytes dies by apoptosis before completion of autosomal synapsis. (A) Bouins-fixed testis sections stained with Periodic acid-Schiff staining. Arrows indicate apoptotic cells in tubule at epithelial stage IV. Post-IV indicates a tubule at a stage past stage IV, where spermatocytes have completely disappeared. Pre-IV indicates a stage before stage IV. (B) Quantification of apoptotic cells in chromosome spread preparations of the indicated genotypes following TUNEL staining of chromosome spreads co-stained with anti-SYCP3 and anti-SYCP1 antibodies (three mice analyzed for each genotype). Error bars = mean±SD; p= p value. (C) Representative images of nuclear spreads of *H2afx^+/+^* and *H2afx ^-/-^* spermatocytes labeled with anti-SYCP3, anti-SYCP1 and TUNEL at zygonema and pachynema stages. Scale bar is 5*μ*m.

### Lack of *H2afx* partially impairs the correct processing of DSBs

Building on the above findings, we hypothesized that *H2afx* null spermatocytes suffered from a defect in the processing of DSBs. In meiotic cells, repair of DSBs is characterized by the timely recruitment and release of several DSB-repair factors (e.g. see (Moens et al., 2002, La Volpe and Barchi, 2012)). Among protein factors involved in early stages of recombination, we determined the kinetics of formation of the Breast Cancer Type 1 susceptibility protein (BRCA1), a homologous-recombination repair protein that we found to assemble into foci in a DSB-dependent manner (Fig. 5A-B). We observed that, in *H2afx^-/-^*mice, foci were present at leptonema and early/mid-zygonema cells, whose quantity was similar to the wild type’s (Fig. 5A and S1 table). This confirms that (as observed in Fig. 1B), lack of H2AFX does not affect DSBs formation and repair at these stages. Interestingly, however, the quantification of DMC1 foci number in late-zygonema cells of *H2afx* mutants revealed, by this stage, an increasing trend in the number of foci. Moreover, the foci number was significantly higher, compared to control, in (H1t-negative) early-pachynema nuclei (Fig. 5C, S1 table). This observation (that was further confirmed in cells from adult mice, see below) suggests a defect in the proper processing of a subset of DSBs. Alternatively, the increase in DMC1 foci number may possibly be caused by the occurrence of a late wave of SPO11-mediated DSBs. However, if *H2afx* was needed to constrain SPO11 activity, we would have expected the DSBs number to also be high at leptonema and zygonema stages. Therefore, we consider the latter hypothesis unlikely. To further understand whether the lack of *H2afx* might impact on later recombination events, we quantified the MSH4 foci number. MSH4 is a member of the mammalian mismatch repair gene family, which binds onto recombination intermediates (i.e. Holliday Junctions), and stabilizes homolog interaction, promoting synapsis (Kneitz et al., 2000, Snowden et al., 2004). In wild type mice, cytologically visible foci of MSH4 appear first in early synapsed regions (i.e. synaptic forks) at early-zygonema (Moens et al., 2007, Storlazzi et al., 2010). Subsequently, their number picks up along synapsed cores at late-zygonema, and a subset of foci persists until mid-pachynema, where discrete MSH4 foci co-localize with late recombination markers, comprising MLH1 (Santucci-Darmanin et al., 2000). We observed that, compared to wild type cells, *H2afx^-/-^* spermatocytes displayed a small but significantly reduced number of foci at both zygotene and early-pachytene stages (Figs 5 D-E, S1 table). This suggests a defect in proper assembly or stable binding of MSH4 onto chromosomes. However, since mice used in this study were in a mix C57/129 genetic background, the observed defect might simply result from small differences in the loading of the protein, due to a random segregation of modifier genes. In order to understand if that was the case, we repeated MSH4 foci count on a different triplet of mice, born from a different breeding pair, at least ten generation away from the previous one. We confirmed that the number of MSH4 foci was significantly smaller (S5 A Fig.).

**Fig.5.**
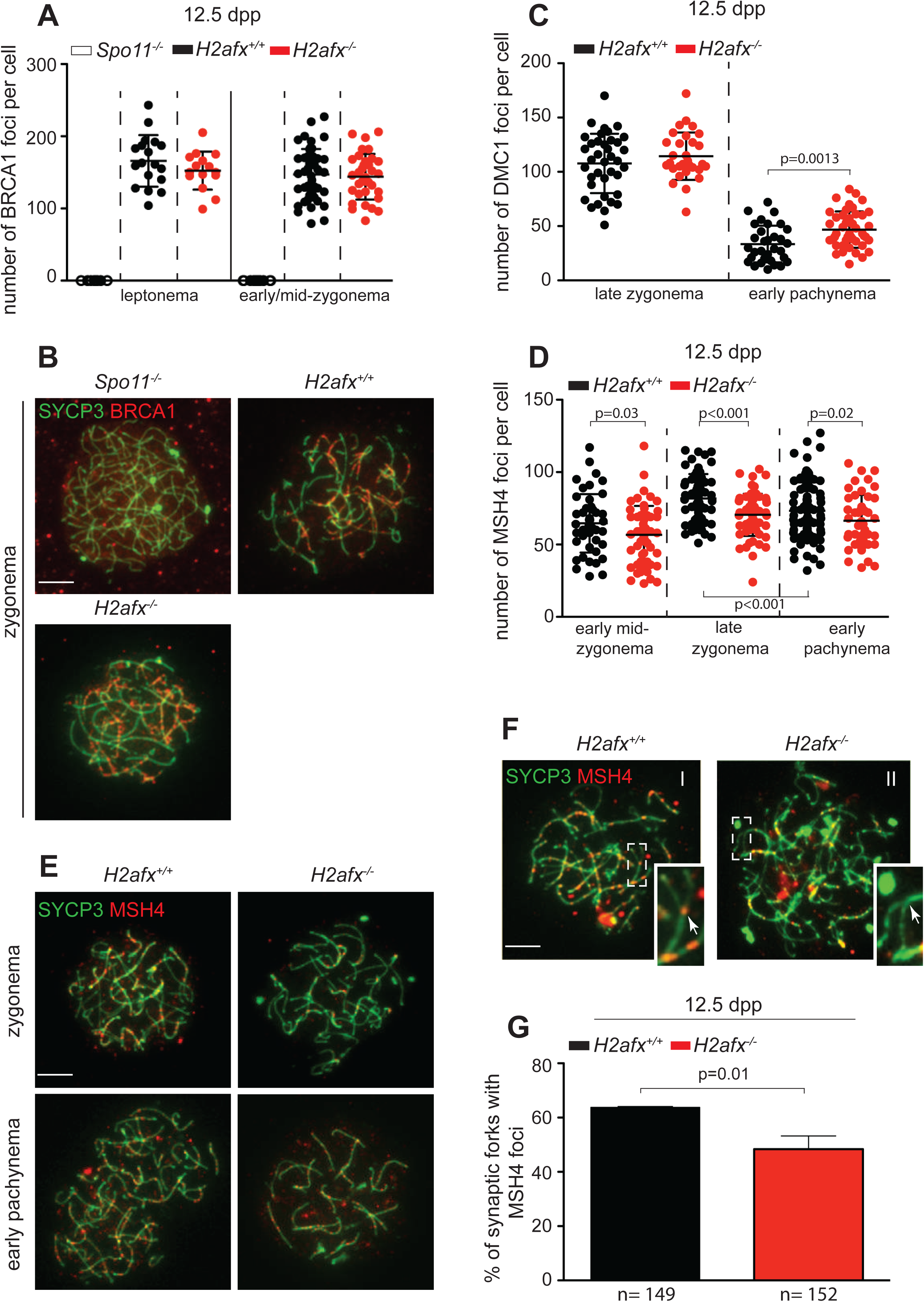
Lack of *H2afx* impairs DMC1 and MSH4 turnover. (A) Quantification of BRCA1 foci number in mice of the indicated genotypes. Each dot in the graph indicates the number of DMC1 foci per nucleus; the error-bars= means ± sd (see S1table for more details). Three mice analyzed for each genotype. (B) Representative images of localization of BRCA1 (red) along SYCP3-positive (green) chromosome axes of spermatocytes at zygonema, in the indicated genotypes. (C) Quantification of DMC1 foci number in the indicated genotypes. Three mice analyzed for each genotype (see S1 table for more details). (D) Quantification of MSH4 foci number in mice of the indicated genotypes. Each dot in the graph 5C and 5D indicates the number of foci per nucleus; error-bars= means ± sd; p= p value. (see S1 table for more details). (E) Representative images of nuclear spreads of indicated genotypes labeled with anti-SYCP3 and anti-MSH4 antibodies. Scale bar represents 5*μ*m. (F) Representative images of nuclear spreads of indicated genotypes labeled with anti-SYCP3 and anti-MSH4 antibodies. Enlarged images are magnifications of dotted areas. The white arrows point to MSH4 foci at representative synaptic forks. Scale bar is 5*μ*m. (G) Quantification of Msh4 foci number in synaptic forks of spermatocytes of the indicated genotypes. (two mice analyzed for each genotype). Error bars = mean±SD; p= p value.

We therefore concluded that lack of *H2afx* impacts on the proper formation of MSH4 foci. The low presence of MSH4 foci at recombination intermediates, might either be due to the premature release of the protein, or to its poor assembly. In this respect, we noticed that while the number of MSH4 foci declined significantly from late-zygonema to early-pachynema in wild-type cells, their number remained substantially unchanged in *H2afx^-/-^* spermatocytes (Fig. 5D, S1 table, S 5A Fig.). This indicates that in the absence of H2AFX, the release of MSH4 from DNA is delayed, pointing to a reduced assembly of MSH4 as the underlying mechanism for the nucleus-wide reduction of foci number. To further test this hypothesis, as MSH4 foci are first cytologically visible at synaptic forks (Fig. 5F I), we quantified fork-associated foci in both wild-type and *H2afx^-/-^* cells. We observed a reduction in MSH4 foci associated with synaptic forks in the mutant (Figs 5 F-G), which confirmed our assumption. Importantly, the number of MSH4 foci in *Mdc1* mutant spermatocytes was similar to that of control (S1 table). This indicates that, contrary to *H2afx, Mdc1* is dispensable for the assembly of MSH4 onto foci. Overall, our findings demonstrated that H2AFX is needed for proper processing of DSBs, probably by implementing the assembly of MSH4 onto DNA.

### Both *H2afx* and *Mdc1* promote deposition of MLH3 foci at pachynema

In the pachytene stage cells of *Mdc1* and *H2afx* null spermatocytes, the assembly of MLH1 foci along the chromosome axes is severely reduced (Ichijima et al., 2011, Celeste et al., 2002). However, females are fertile (Celeste et al., 2002, Lou et al., 2006). These observations, along with the fact that in normal males MLH1 appears by the time cells of the mutants undergo apoptosis (Kolas et al., 2005, de Boer et al., 2006, Moens et al., 2007, Mahadevaiah et al., 2008, Ahmed and de Rooij, 2009), has led to the conclusion that the defective assembly of MLH1 in males reflects the fact that cell death precedes foci formation. However, an alternative interpretation is that the altered deposition of MLH1 reflects a problem in the execution of the pro-crossover pathway. To distinguish between these alternative interpretations, we stained wild-type, *H2afx^-/-^* and *Mdc1^-/-^* mice spermatocytes with an antibody that recognizes the mismatch-repair protein 3 (MLH3), a pro-crossover factor that appears at early pachynema, before cells of the mutants undergo apoptosis at stage IV (Lipkin et al., 2002, Kolas et al., 2005, Svetlanov et al., 2008). Quantification of MLH3 foci in both *H2afx^-/-^* and *Mdc1^-/-^* cells revealed that, although MLH3 foci were present and increased significantly during the early pachynema (H1t negative) to early mid-pachynema (faintly H1t positive) transition, their global number was significantly smaller, relatively to control (Figs 6 A-C). This observation indicates that *H2afx* and *Mdc1* are needed to promote the assembly or stable binding of MLH3 onto DNA. However, whether this phenotype is the consequence of either a delay or an arrest of the pro-crossover pathway in males, remains to be elucidated (see discussion).

**Fig.6.**
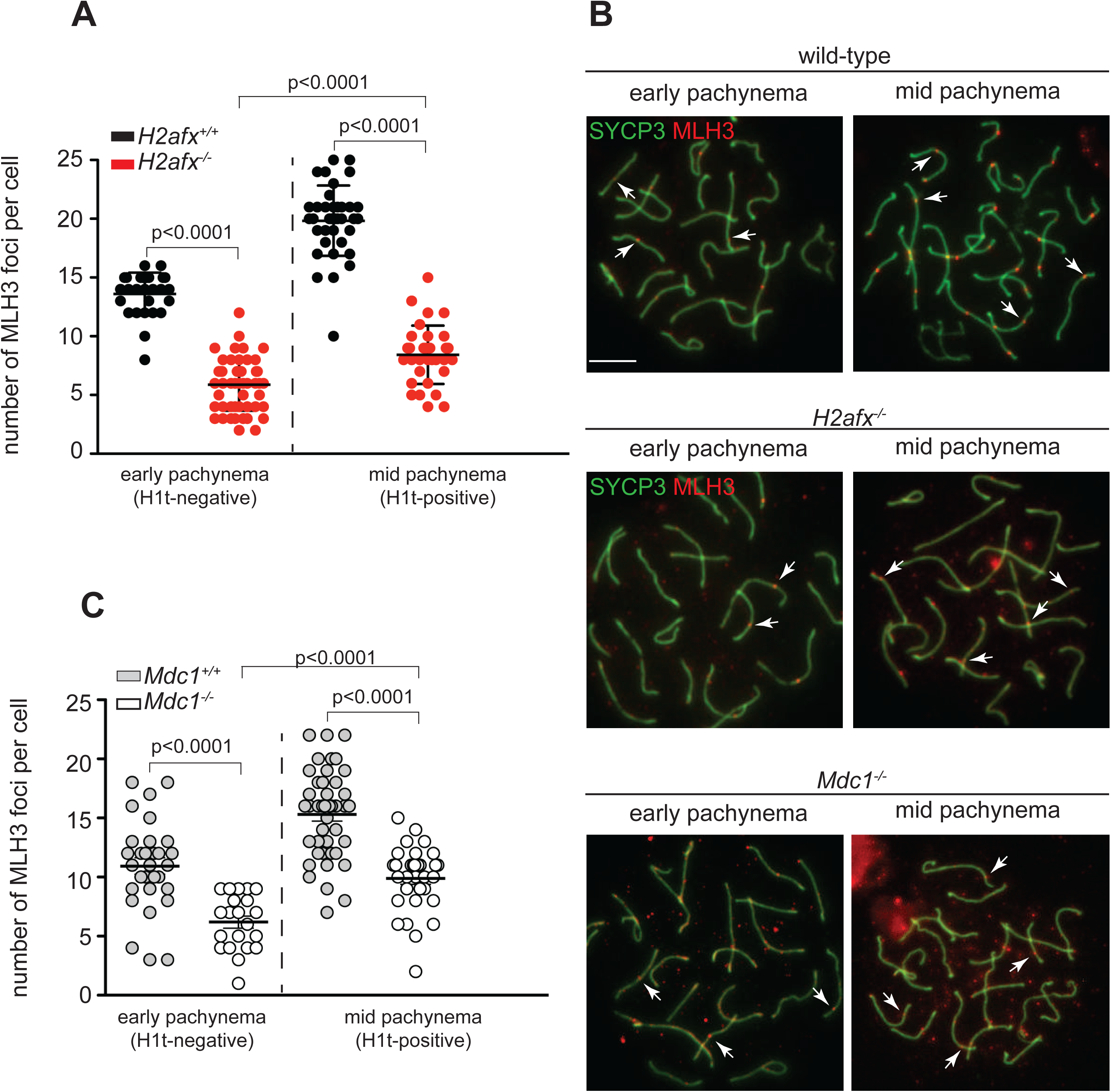
*H2afx* and *Mdc1* mutant spermatocytes are defective in MLH3 foci assembly. (A and C) Quantification of MLH3 foci in 60 dpp mice of the indicated genotypes and stages (three mice analyzed for each genotype). Each dot in the graph indicates the number of MLH3 foci per nucleus; the black bars are means and standard deviation, p= p values (see S1 table for more details). (B) Representative images of chromosome spreads from the indicated stages and genotypes, stained with antibodies against SYCP3 (green) and MLH3 (red). White arrows point MLH3 foci.

### *H2afx* and *Mdc1* tie resolution of DSBs with progression of spermatocytes to mid-pachynema

In mouse spermatocytes, mutations that leads to an increase in the number of DSBs at early pachynema, or prevent DSBs from being repaired, causes a delay or arrest of spermatogenic differentiation at early pachynema, following the activation of the recombination-dependent checkpoint (Pacheco et al., 2015, James H Crichton, 2017). Accordingly, in mutants where the checkpoint is bypassed, normal co-ordination of progression with resolution of DSBs is altered, and cells enter mid-pachynema (i.e. incorporate the mid-pachytene stage marker H1t); besides, DNA damage persists. The activation of the recombination-dependent checkpoint in mice, relies on the function of ATM (Pacheco et al., 2015). Since H2AFX is a phosphorylation target of ATM (Rogakou et al., 1998, Barchi et al., 2008), and in somatic cells H2AFX phosphorylation is required to reinforce ATM signaling (Lou et al., 2006), we reasoned that H2AFX might be involved in promoting the activation of the recombination-dependent checkpoint. To determine whether lack of *H2afx* affects co-ordination between meiotic progression (i.e. incorporation of H1t) and repair of DSBs at pachynema, we used indirect immunofluorescence with antibodies against SYCP3, DMC1 and H1t, on spermatocytes chromosome preparations. At first (as we observed in 12.5 dpp *H2afx^-/-^* mice, Fig. 5C), by analyzing the cells of adult mice, we confirmed our observation that H1t-negative pachytene spermatocytes showed higher numbers of DMC1 foci (Fig. 7A and 7B I-II). In addition, we found that H1t-positive *H2afx^-/-^* spermatocytes also had numerous DMC1 foci (Fig 7A and 7B III-IV), only ~ 28% fewer than H1t-negative *H2afx^-/-^* cells. This was remarkably different from wild type cells, were most of the cells that became H1t-positive had much lower number of DSBs (~ 83% of reduction) than the early H1t-negative cells (Figs 7A and 7B I and III). Moreover, H1t-positive *H2afx^-/-^* spermatocytes had significantly more DMC1 foci than H1t-positive wild type cells (Fig. 7A). This suggests that, in absence of *H2afx*, spermatocytes progress to mid-pachynema, despite the persistence of a high number of DSBs. If this hypothesis is correct, as observed in other mice models (Pacheco et al., 2015), we would also expect to observe an increase in the percentage of pachytene stage cells that progress to mid-pachynema, relatively to wild type. We thus examined H1t incorporation in spermatocytes of wild type and *H2afx^-/-^* at 12.5 dpp old mice. In wild type, 35±5% of pachytene stage cells were positive for H1t, while the percentage increased significantly in *H2afx^-/-^* testes (54± 2%, p<0.05) (Fig. 7C). It is worth highlighting that, in H1t-positive cells of 12.5 dpp spermatocytes in the *H2afx* mutant, the number of DMC1 foci was significantly higher relatively to wild type (S 5B Fig.), thus confirming the result obtained using adult mice (Fig. 7A). Overall, these observations indicate that *H2afx* mutant spermatocytes have a defect in coordinating repair of DSBs with meiotic progression. In order to understand whether *Mdc1* is functionally linked with *H2afx* in this function, we quantified the number of DMC1 foci in different substages of pachynema in *Mdc1^-/-^* spermatocytes from adult mice. We observed that, at early pachynema, the average number of DMC1 foci was not different from that of control (Fig. 7D). Nevertheless, while the number of DSBs in cells that progressed to midpachynema was smaller by ~ 70% in wild type, such decrease was much milder (~ 30%) in *Mdc1* mutant cells (Fig. 7B VII-IX and 7D). Since in *Mdc1^-/-^* spermatocytes the dynamic of DSB formation and repair was normal up to early pachynema, the observed increase in DMC1 foci in H1t-positive cells is unlikely to be due to a late wave of DSBs, or to a defect in DSB repair. Therefore, we inferred from this result that *Mdc1* is needed to impose a delay at early-pachynema, which prevents premature progression to mid-pachynema. Accordingly, the count of H1t-positive cells in 12.5 dpp testes revealed that, in the mutant, the number of H1t-positive cells had increased (wt: 23±2%; *Mdc1^-/-^*: 37%±1, p<0.05) (Fig. 7E), and in H1t-positive cells, the number of DMC1 foci was higher relatively to wild-type cells (S 5C Fig.) confirming data obtained with adult mice.

**Fig.7.**
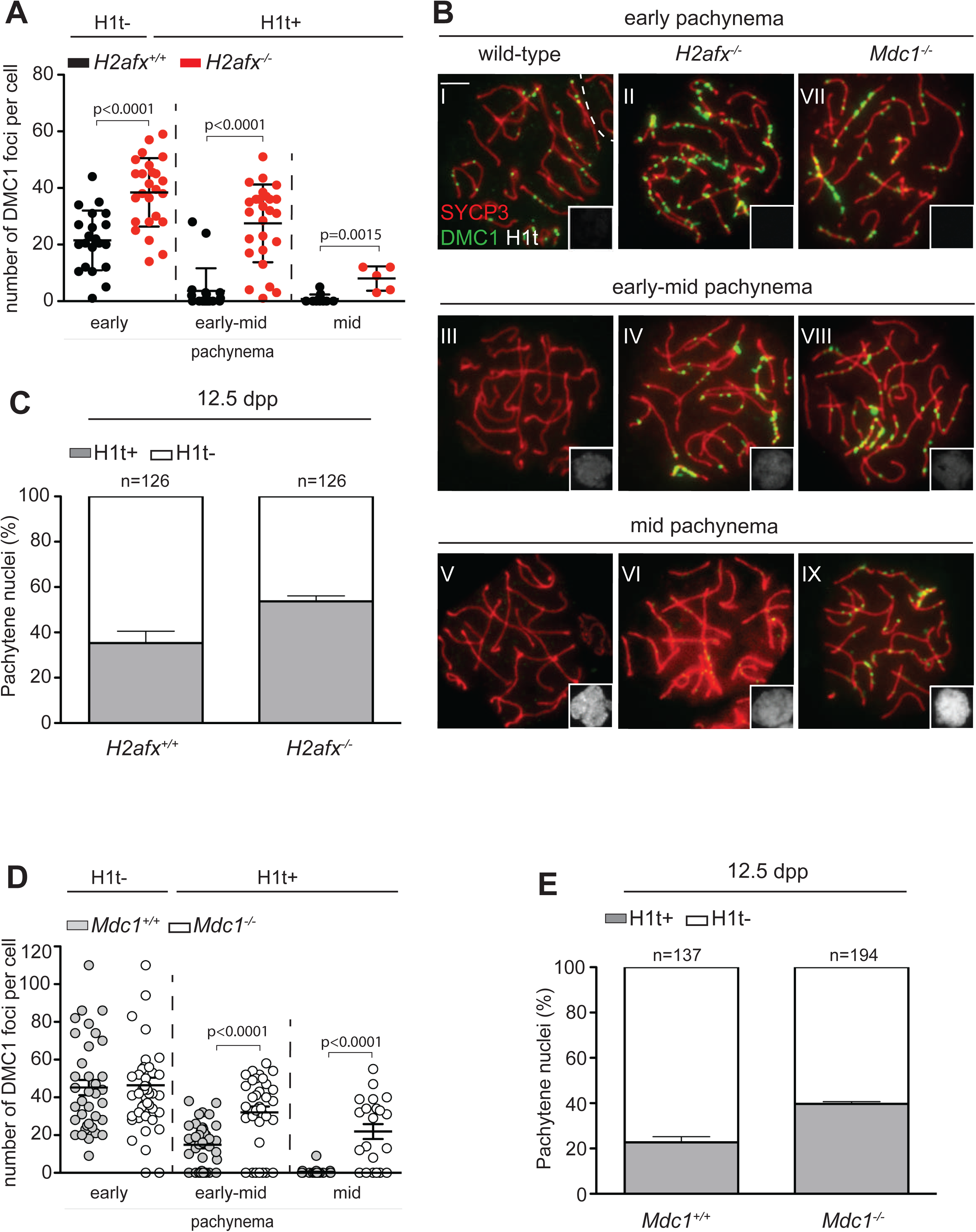
*H2afx* and *Mdc1* genes tie DSBs repair with meiotic progression. (A and D) Quantification of DMC1 foci number in adult mice of the indicated genotypes and stages. Sub-classes of spermatocytes were identified by looking at the intensity of H1t staining (namely: no staining= early pachynema, faint staining= early to mid-pachynema, intense staining= mid-pachynema). Each dot in the graphs indicate the number of DMC1 foci per nucleus. Three mice analyzed for each genotype. (B) Representative images of nuclear spreads of indicated genotypes labeled with anti-SYCP3 (red), anti-H1t (white, in the insert) and anti-DMC1 (green) in cells with the indicated genotypes. Magnification bar is 5*μ*m. (C and E) Percentage of H1t-positive and H1t-negative spermatocytes from mice of the indicated genotypes. The black bars in A and D are means and SD. p= p values. Error bars in C and E are SD. n= number of analysed cells.

It is important to stress that, the phenotypes observed in both mutants are unlikely to be linked to the over-expression of H1t, as transcriptional levels were found to be comparable to that of wild type (S6 A-B Figs.). In addition, comparing the timing of H1t expression in testis sections from wild type, *H2afx* and *Mdc1* mutants, we found that, as expected (Mahadevaiah et al., 2008), in controls, H1t was first detected in mid-pachytene spermatocytes at stage IV (S6 C Fig., top panel). Staging is less accurate for mutants that lack post meiotic cells. Nonetheless, in most cases, H1t-positive cells were found in stage IV tubules (i.e. containing numerous apoptotic spermatocytes (Mahadevaiah et al., 2008, Ahmed and de Rooij, 2009)), (S6 C Fig., mid and lower panels), which is against the possibility that *H2afx* and *Mdc1* are required to prevent premature loading onto the chromatin of H1t. Altogether, our observations suggested that, in mouse spermatocytes, *H2afx* and *Mdc1* are needed to tie the resolution of DNA damage with meiotic progression through prophase I, a phenotype reminiscent of that of mutants with a defect in the activation of the recombination-dependent checkpoint (Pacheco et al., 2015, James H Crichton, 2017, Marcet-Ortega et al., 2017).

### *Dmc1^-/-^ H2afx^-/-^* spermatocytes progress to an H1t-positive state in greater numbers, relatively to *Dmc1^-/-^*

To further challenge the hypothesis of a role for *H2afx* in constraining progression of spermatocytes with unrepaired damage, we sought to compare H1t incorporation in *Dmc1^-/-^* and *Dmc1^-/-^ H2afx^-/-^* double mutants. *Dmc1* deficiency prevents synapsis and DSBs repair (Pittman et al., 1998), and spermatocytes were reported not to incorporate H1t (Barchi et al., 2005, Pacheco et al., 2015), or incorporate it at low level (Mahadevaiah et al., 2008). In line with previous results, using chromosome spreads analyses, we observed that only a small portion of *Dmc1^-/-^* spermatocytes was faintly H1t-positive. This percentage increased significantly in *Dmc1^-/-^H2afx^-/-^* double mutants (Fig. 8 A-B) (*Dmc1^-/-^* 25.7±2%; *Dmc1^-/-^H2afx^-/-^*47.6±8%), which was suggestive of a less penetrant arrest or delay of progression in the double mutant. In spermatocytes, the incorporation of H1t is not prevented by a simple defect of chromosome synapsis (i.e. as if the lack *Spo11* ((Barchi et al., 2005, Pacheco et al., 2015)). Hence, our data represents a direct indication that *H2afx* supports the activation of a mechanism that constrains the meiotic progression of cells with unrepaired damage. Interestingly, by comparing the intensity of H1t signal in *Dmc1^-/-^* and *Dmc1^-/-^H2afx^-/-^* side-by-side, we observed that, in most cells, the H1t incorporation levels were comparably low (S 7C Fig.), as if the cells did not progress over early-mid pachynema. Conversely, in both *H2afx^-/-^* mutant cells (Fig. 7B VI), and in spermatocytes where the recombination-dependent checkpoint is bypassed at pachynema (Pacheco et al., 2015, Marcet-Ortega et al., 2017), nuclei progressed up to mid/late pachytene stage. This suggested that, in pachytene-like cells of *Dmc1^-/-^* spermatocytes, lack of *H2afx* only promotes partial release of the checkpoint that restrains their progression to mid pachynema. Therefore, additional mechanisms are likely to be in place, which are independent from the function of *H2afx* (see discussion).

**Fig. 8.**
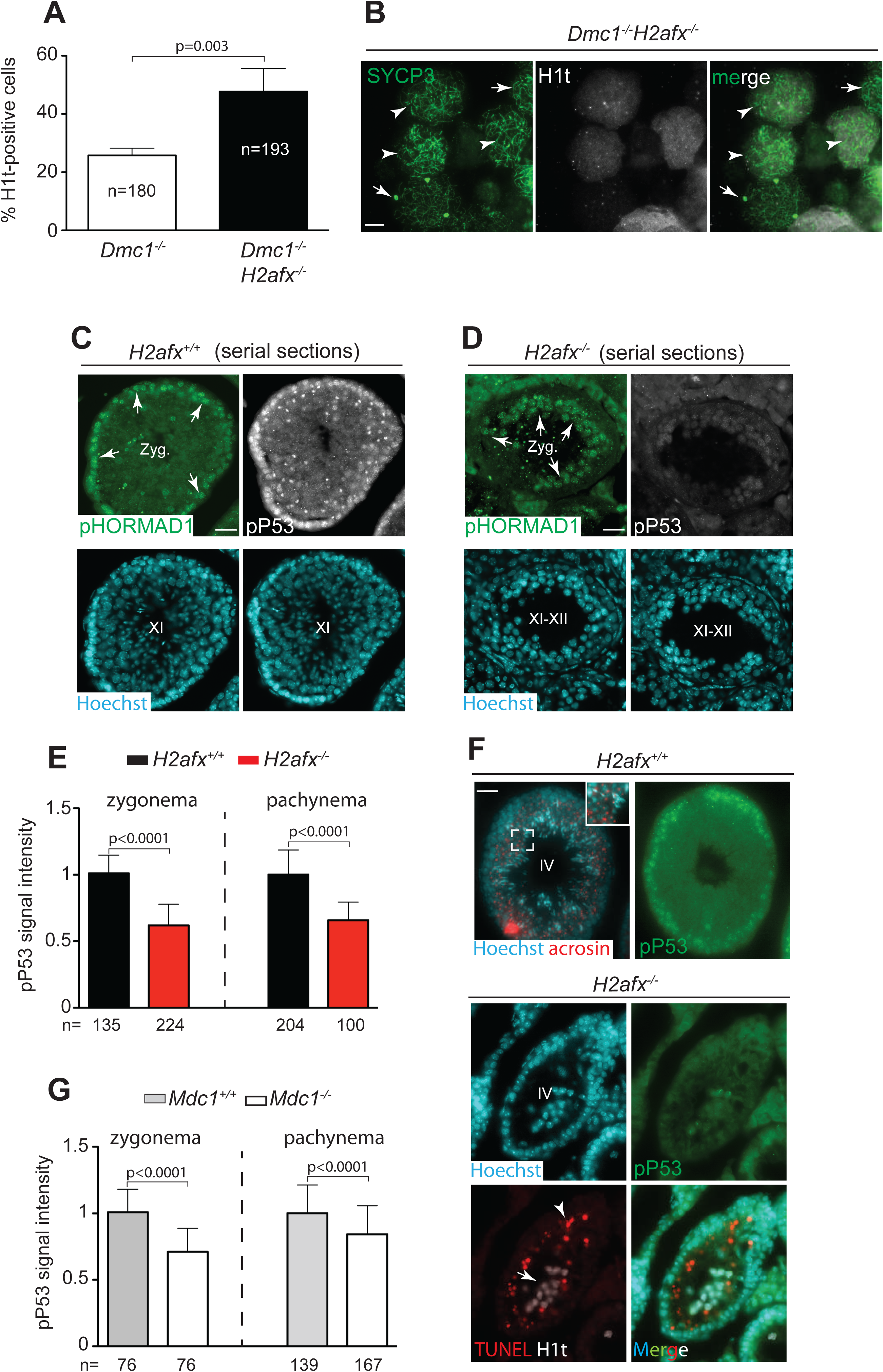
*H2afx* supports the activation of the recombination-dependent arrest and along with MDC1 promotes the DSB-mediated activation of P53. (A) Quantification of H1t-positive cells from the indicated genotypes. (B) Representative image of H1t staining signal in *Dmc1^-/-^H2afx^-/-^* spermatocytes. Arrows head point to H1t-positive cells, while the arrows point to H1t-negative nuclei. Scale bar= 5*μ*m. (C-D) Representative images of immunofluorescence staining of serial sections from testes of the indicated genotypes, stained with the anti-pP53 and anti phspho-HORMAD1^375^ (pHORMAD1) antibodies. White arrows indicate spermatocytes in the zygotene stage (Zyg.). Roman numbers indicate the stages of the epithelial cell cycle. (E) Quantification of pP53 signal in spermatocytes at the indicated stages of development, in wild-type and *H2afx^-/-^* mice. (F) Top panel; representative image of a wild-type seminiferous tubule at stage IV of the epithelial cell cycle, stained with Hoechst (blue), acrosin (red) and pP53 (green). The enlargment shows the pattern of the anti-acrosin antibody in round spermatids. Lower panel; representative image of a stage IV tubule in *H2afx^-/-^* testis, stained with Hoechst (blue), pP53 (green), TUNEL (red) and H1t (white). The arrow point H1t-positive cells, while the arrow head indicate apoptotic spermatocytes. (G) Quantification of pP53 signal in spermatocytes at the indicated stages of development, in wild-type and *Mdc1^-/-^* mice. Magnification bars in C, D and F are 25*μ*m. Error bars in A, E and G are SD, n= number of analysed cells.

### Deletion of *H2afx* and *Mdc1* dampens P53-mediated activation to DSBs

In mouse spermatocytes, activation of the recombination-dependent arrest at pachynema relies on the function of P53 (Marcet-Ortega et al., 2017). We reasoned that, if *H2afx* and *Mdc1* are required to support the recombination-dependent arrest in spermatocytes, then they might be required to promote the activation of P53. In mouse testis, Spo11-mediated P53 activation can be monitored by looking at its phosphorylation on Ser^15^ (pP53) (Lu et al., 2010). By immunostaining testes sections with the anti-pP53 antibody, and on the basis of nuclear morphology and of the staining pattern of the anti-acrosin antibody (which stains the developing acrosome (Muciaccia et al., 2013)), we characterized the pP53 pattern in developing spermatocytes, in wild-type. From leptonema to early pachynema, pP53 appeared as diffuse staining through the nucleus (S7 A i and S7 A iv-v Figs). At subsequent substages, it becames gradually localized in a single nucleus area (S7 A ii-iv Figs), and it disappeared at diplonema (S7 A v Fig.). In those experiments, the pP53 signal was specific, because it was absent in testes sections from *p53^-/-^* mice (S 8A Fig.). Chromosome spreads analysis further revealed that at zygonema, a fraction of pP53 in the nucleus was localized on the chromosome axes of unsynapsed chromosomes (S7 B I Fig.). When cells progressed to early pachynema, chromatin-bound pP53 became restricted to the axes of unsynapsed sex chromosomes (S7 B II Fig.), and extended over chromatin loops of X-Y chromosomes by mid-pachynema (S7 B III Fig.).

In somatic cells, P53 is a target of the ATM/Chk2 DNA-damage response pathway (Canman et al., 1998, Khanna et al., 1998, Banin et al., 1998). With a view to verifying that, in spermatocytes, ATM and P53 are in the same pathway, we stained testes sections of *Atm^-/-^* mice with the anti-γH2afx and -pP53 antibodies. We focused on cells at leptonema, because as demonstrated by the lack of γH2afx staining, at that stage, the ATM function is not likely replaced by ATR (Bellani et al., 2005, Barchi et al., 2008, Royo et al., 2013). We observed that, in γH2afx –negative (i.e. leptotene stage (Bellani et al., 2005, Barchi et al., 2008, Royo et al., 2013)) cells, pP53 was not apparent (compare Fig. S 8 B and S 8 C). This confirmed that P53 is a target of ATM. It is interesting to highlight that, in *Atm^-/-^* spermatocytes where γH2afx signal was present (i.e. zygotene stage nuclei (Bellani et al., 2005, Barchi et al., 2008, Royo et al., 2013)), the pP53 staining was also often visible, albeit with often reduced intensity relatively to wild-type (Fig. S 8 C). This suggests that, at least in this genetic context, P53 is also a target of ATR.

ATM signaling is canonically enhanced by a positive-feedback loop mechanism that involves MDC1 binding to γH2AFX (Yuan et al., 2010, Coster and Goldberg, 2010). To investigate whether H2AFX and MDC1 are components of the ATM>P53 signaling cascade, we tested whether they are required to support P53 phosphorylation. By measuring the pP53 signal intensity in testis sections of *H2afx^-/-^* and *Mdc1^-/-^* mice, we observed that, in pHORMAD1-positive zygonema cells, the P53 phosphorylation signal was dampened, with a reduction of about 30% in *Mdc1* mutant and 43% in *H2afx^-/-^* spermatocytes (Fig. 8C-E and 8G). To further test whether pP53 was also reduced in cells that progressed to early-mid pachynema, we quantified it in spermatocytes at stage IV of the epithelial cell cycle. In wild-type, spermatocytes at stage IV were recognized by staining testes sections with the anti-acrosin antibody (Fig. 8F, top panel [enlargement]). In *H2afx^-/-^* and *Mdc1^-/-^* mice, spermatocytes were almost completely eliminated by apoptosis at stage IV of the epithelial cell cycle (Mahadevaiah et al., 2008, Ahmed and de Rooij, 2009). We therefore measured pP53 in TUNEL-negative/H1t-positive cells, in sections of seminal tubules that contained numerous apoptotic cells (Fig. 8F, lower panel). The analyses revealed that, in TUNEL-negative/H1t-positive spermatocytes of *H2afx^-/-^* and *Mdc1^-/-^* mice, pP53 was reduced, relatively to wild-type, to almost the same extent as that of cells at zygonema (Fig. 8E and 8G). It is worth highlighting that, the weakening of the pP53 signal was not due to poorer expression of P53 in absence of *H2afx* or *Mdc1*, as P53 protein was found to be present at similar levels to control ones (S8 C Fig.). We concluded that, in spermatocytes, H2AFX and MDC1 support the activation of P53, in response to Spo11-mediated DSBs.

## Discussion

In all sexually reproducing organisms, DSBs are formed as part of the normal developmental program. Indeed, the formation and repair of DSBs through homologous recombination leads to the formation of COs, the physical link that is essential for correct chromosome segregation during prophase I. In mammals, as well as in all organisms that additionally rely on recombination for efficient pairing of the homologs (Bolcun-Filas and Schimenti, 2012), formation of SPO11-mediated DSB is massive (Keeney, 2008). This creates, in meiotic cells, opportunities to generate gross chromosomal rearrangements (Kim et al., 2016). As a consequence, germ cells are expected to possess mechanisms that control the accurate performance of the recombination process, and delay or arrest their progression through differentiation, if recombination is defective. Such behavior prevents genomic instability, with the generation of a healthy offspring. In our study, we uncovered the role of *H2afx* in promoting proper meiotic recombination, which is reminiscent of the function of *H2afx* in somatic cells (Xie et al., 2004, Sonoda et al., 2007). By using the MLH3 recombination marker, we also demonstrated that both *H2afx* and *Mdc1* are required for the proper execution of the pro-crossover pathway, that leads to CO formation. Lastly, we showed evidence that *H2afx* and *Mdc1* are required to delay or arrest the progression through differentiation of pachynema cells with persistent DNA damage. Overall, our results suggest that *H2afx* and *Mdc1* play a greater role in preventing genome instability in meiotic cells, than it was previously apparent.

### H2AFX enforces recombination-mediated synapsis of the autosomes

In somatic cells, besides fulfilling other functions, H2AFX and MDC1 promote the maintenance of genome stability, by enforcing homologous recombination-mediated DSB repair (Unal et al., 2004, Xie et al., 2004, Sonoda et al., 2007, Bassing et al., 2002, Xie et al., 2007, Xie et al., 2010). However, whether these proteins also play such a role in meiosis is unclear. Previous findings had shown that, in *H2afx* mutant, synapsis of the autosomes is defective (Moens et al., 2007). Whether the observed problem is due to a defect in DSBs formation or to their correct processing, remained unknown. By analyzing DMC1 deposition on bulk chromatin, we found that, in the absence of *H2afx*, DSBs form in normal numbers. This is consistent with the observed normal localization of BRCA1 and levels of phosphorylation of HORMAD and SMC3 proteins (Fukuda et al., 2012). Despite this, chromosome dynamic was altered in a significant fraction of *H2afx* mutant cells at zygonema, thus suggesting that there was a defect in recombination. In support of this hypothesis, we also observed that, similar to mouse mutants where recombination is compromised (Barchi et al., 2005), a relevant fraction of cells that had no completed synapsis died by apoptosis at stage IV of the epithelial cell cycle. This was in sharp contrast to what we observed in *Mdc1^-/-^* mice, which did not show any relevant defect in timely progression of recombination and chromosome synapsis. We interpreted this result to mean that a fraction of cells at zygonema failed to complete synapsis of the autosomes in a timely manner. It is important to stress that, since the nuclei of most TUNEL-positive cells were too shrunken to be informative on the stage of cell death, the estimated percentage of cells with delayed synapsis is likely to have been underestimated. These observations point to a much more significant role for *H2afx* in controlling appropriate and timely synapsis of the autosomes, compared to what was previously anticipated. We recently demonstrated that, in a mouse mutant with no defects in DSB processing, zygonema-like cells that receive late surge DSBs are still able to complete chromosome synapsis on time, with no overt increase of apoptosis (Faieta et al., 2015). This observation strengthens the hypothesis that *H2afx* is required to enforce meiotic recombination. In line with with the above results, we observed that the number of DMC1 foci had significantly increased in mutant cells at early pachynema. In addition, in *H2afx^-/-^* nuclei the number of MSH4 foci was significantly lower at both zygotene and pachytene stages. The MutSγ complex, represented by MSH4 and MSH5, play in mammals (as well as in many other organisms), a well-established role in the stabilization of the Holliday Junctions intermediates and homologous synapsis (Edelmann et al., 1999, Kneitz et al., 2000, Snowden et al., 2004, Kolas et al., 2005). From these observations we inferred that H2AFX promotes synapsis of the autosomes, by enforcing recombination proficiency. As H2AFX is a histone, we speculate that it might contribute to shape chromatin conformation and accessibility to DNA repair factors, including MSH4 (Fig. 9). This hypothetical function may depend on presence of H2AFX *per se*, or on the absence of some H2AFX post-translational modifications induced by DNA-damage (Xie et al., 2010). The analysis *in vivo* would require the generation of mice models carrying mutations at specific post-translational target sites. On the other hand, since in somatic cells H2AFX-mediated function in recombination largely relies on its phosphorylation (Sonoda et al., 2007), we suppose that γH2AFX might be required to support meiotic recombination. This interpretation was corroborated by the observation that, proper repair of DSBs in *Mdc1* mutant spermatocytes correlated with the presence of (albeit reduced) γH2AFX signal along chromosome axes (Lee et al., 2005). This interpretation is also consistent with the findings relating to somatic cells, where limited formation of γH2AFX, in absence of *Atm*, is sufficient to support *H2afx*-dependent homologous recombination (Rass et al., 2013).

**Fig. 9.**
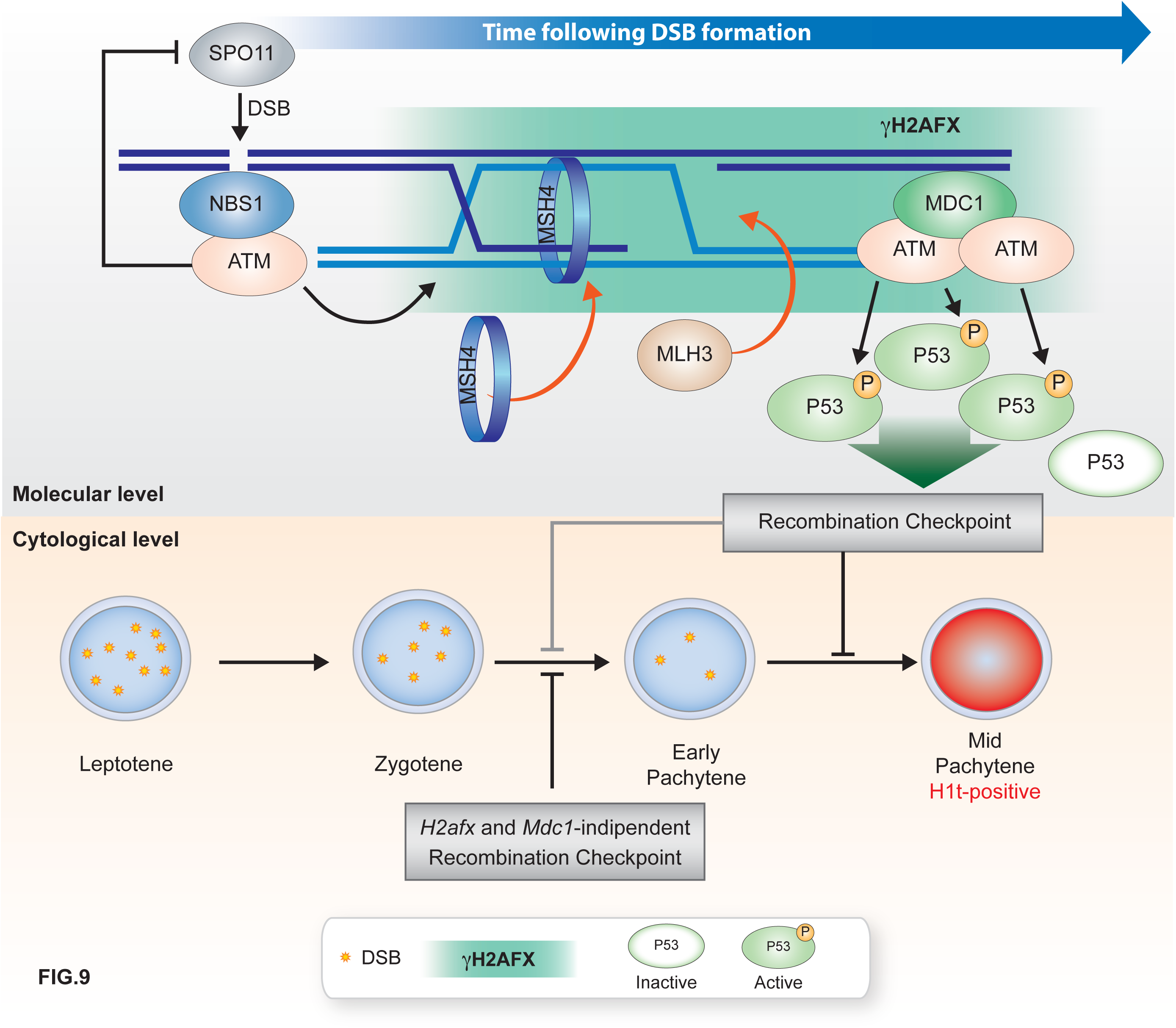
Schematic model describing H2AFX and MDC1 functions in response to Spo11-mediated DSBs, in spermatocytes. Based on the information present in this and previous studies we propose that following DSBs formation, activated ATM operates a feedback control on Spo11 catalytic activity (Lange et al., 2011) which does not require *H2afx* or *Mdc1* functions (black line, top panel). Following the phosphorylation of H2AFX (γH2AFX) by ATM, chromatin conformation might be modified to assist proper recruitment of MSH4 at ongoing recombination sites. This supports stable alignment and pairing of the homologs, ensuring that synapsis occurs properly and is timely completed, at zygonema. When cells reach early-pachynema, MLH3 is recruited, or it is stabilized onto chromatin, possibly through the interaction with a factor(s) that is loaded onto chromatin, with a mechanism that rely on the presence or post-translational modification(s) of H2AFX, with the contribution of MDC1. Concomitantly, the MDC1-mediated recruitment of ATM on γH2AFX, amplify the ATM response to damage, including the activation of P53 by its phosphorylation on Ser^15^ (active P53). Phosphorylated P53 activates the recombination-dependent checkpoint that partially constrain progression of cells at zygonema (light grey line, lower panel), and prevents untimely progression of early-pachynema cells with eccessive unrepaired DSBs, to mid pachynema (black line, lower panel) (Marcet-Ortega et al., 2017). DSB= double strand break.

### *H2afx* and *Mdc1* play partially overlapping functions in X-Y synapsis

Previous findings had shown that both *H2afx^-/-^* and *Mdc1^-/-^* spermatocytes suffered from a defect in X-Y chromosome synapsis (Fernandez-Capetillo et al., 2003, Celeste et al., 2003, Ichijima et al., 2011). However, the relative contribution of those genes to the process was not investigated. We observed that the percentage of X-Y asynapsis in *H2afx^-/-^* spermatocytes was three-fold higher compared to that of *Mdc1^-/-^*. This indicates that *H2afx* and *Mdc1* promote sex chromosomes synapsis through mechanisms that are at least partially distinct. The above observations on synapsis and recombination of the autosomes suggest that sex chromosome asynapsis in *H2afx* spermatocytes is due to the reduced proficiency of recombination, which it was not however observed in mutants lacking *Mdc1*. A common feature of *H2afx* and *Mdc1* mutant spermatocytes is lack of formation of sex body, a chromatin domain that is thought to promote X-Y proximity, by assisting recombination-mediated synapsis at the PAR (Fernandez-Capetillo et al., 2003, Celeste et al., 2003, Ichijima et al., 2011, Kauppi et al., 2011). Therefore, a reasonable interpretation is that X-Y asynapsis in *Mdc1* mutant is due to a reduced chance of sex chromosomes to be spatially close at late zygonema/early pachynema transition, the time when X-Y recombination occurs in normal mice (Kauppi et al., 2011). In *H2afx* mutant spermatocytes, the defect in the formation of the sex body would be additional to that of recombination, resulting in a greater frequency of asynapsis. Therefore, we conclude that, in mice, efficient synapsis of the sex chromosomes depends on both *H2afx* and *Mdc1*, although with only partially overlapping mechanisms.

### H2AFX and MDC1 promote the timely execution of the pro-crossover pathway

In normal cells, out of about 200–300 DSBs that form during early prophase of meiosis I in mouse spermatocytes, only a small subset becomes COs (~20-30). Designation of prospective COs is implemented gradually during meiosis (Cole et al., 2012). At zygonema, the localization of the meiosis-specific MutSγ heterodimer (MSH4/MSH5) to a subset of the initial DSBs reduces the pool of potential CO intermediates by half (Kneitz et al., 2000). The number of MutSγ foci subsequently decline as spermatocytes progress through pachynema, during which time MutLγ heterodimer (MLH3/MLH1) is recruited to a subset of such sites at a frequency and with distribution corresponding to that of the final CO sites (Santucci-Darmanin et al., 2000). MSH4 directly binds MLH3 and MLH1 (Santucci-Darmanin et al., 2002, Santucci-Darmanin et al., 2000) and co-localizes with MLH1 (Santucci-Darmanin et al., 2000), which is likely to promote selective stabilization of pre-CO intermediates (Kolas and Cohen, 2004, Kolas et al., 2005, Cole et al., 2012, Holloway et al., 2008).

In our analyses, we observed that *H2afx* mutant spermatocytes showed a significant reduction in the number of MSH4 and MLH3 foci at pachynema. This indicates an alteration of the pro-crossover pathway. Conversely, while in *Mdc1^-/-^* spermatocytes the number of MSH4 foci was identical to that of control, MLH3 foci were reduced to extent number similar to that of *H2afx* knockout cells. This indicates that the reduction of MSH4 observed in *H2afx^-/-^* cells was not the cause of the alteration of the pro-crossover pathway. It is thus conceivable that *H2afx* and *Mdc1* genes, by contributing to properly shape chromatin, might promote CO formation, perhaps stabilizing MLH3 binding onto DNA, or by favoring the interaction of DNA with factors (such as ZMM proteins other than MSH4 (Lynn et al., 2007, Adelman and Petrini, 2008, Reynolds et al., 2013)) required for the CO designation process, that assemble onto chromosomes in advance of MLH3 (Fig. 9). MDC1 may serve, *per se*, as a scaffold for the recruitment of such factors. Alternatively, as H2AFX is partially phosphorylated in *Mdc1^-/-^* cells, correct MLH3 deposition might require robust levels of (MDC1-mediated) γH2AFX.

One observation against a mechanistic role of *H2afx* and *Mdc1* in CO formation is that, in females, MLH1 foci assemble normally (Celeste et al., 2002, Ichijima et al., 2011). This might depend on the different type and timing of the arrest of defective oocytes. In females, given their X-X complement, MSCI is inactive. Furthermore, the DNA damage-independent and dependent arrests only act in oocytes at late prophase, relatively to males, around the time they enter meiotic arrest (Bolcun-Filas et al., 2014, Cloutier et al., 2015). Given the above-mentioned differences, a possible explanation for the apparent sexual dimorphism in MLH3 and MLH1 foci assembly in *H2afx* and *Mdc1* mutants is that CO designation is delayed in both sexes rather than being arrested. This would allow oocytes from *H2afx* mice to complete synapsis and place (in both *H2afx^-/-^* and *Mdc1^-/-^* genetic backgrounds) MLH3 (and MLH1) at CO sites, with consequently no overt defect in female fertility. In line with this hypothesis, we observed that in males, the average number of MLH3 foci increased through early pachynema to mid-pachynema transition, although they were less numerous compared to control. This indicates that MLH3 foci continued to be deposited, over time, with slow kinetic, before their apoptotic elimination at stage IV of the epithelial cell cycle.

### Arrest of progression prior H1t incorporation at pachynema, likely depends on levels of γH2AFX and pP53

In recombination defective mutants, persistent unrepaired DSBs at pachynema are marked by γH2AFX. The progression of those cells through differentiation (monitored by looking at H1t incorporation) correlates with the lack or lower levels of γH2AFX along autosomes axes, such as in *Trip13* mutant (Pacheco et al., 2015). On the basis of this observation, we argued that in spermatocytes, γH2AFX might represent a signal to constrain cell progression through differentiation. In line with this hypothesis, we observed that in *H2afx^-/-^* mice, the percentage of H1t-positive cells at pachynema was significantly higher compared to wild type, and the number of DSBs in H1t-positive mutant cells was also significantly higher than the control’s. Hence, (as observed for other proteins involved in the ATM signaling pathway (Pacheco et al., 2015)) H2AFX is likely required to implement co-ordination of DSBs repair with meiotic progression. Likewise, although *Mdc1* is dispensable for timely repair of DSBs, the percentage of H1t-positive cells in the mutant was also higher than the wild type’s, and H1t-positive *Mdc1^-/-^* spermatocytes displayed a significantly higher number of unrepaired DSBs compared to its control. This represented the first experimental evidence of the fact that recombination-dependent arrest prior to H1t incorporation is part of the normal developmental program, as it occurs in recombination proficient cells, preventing their premature progression to mid-pachynema. In *Mdc1^-/-^* spermatocytes, H2AFX is phosphorylated at low levels during meiosis (Lee et al., 2005). Therefore, the premature progression of early pachynema cells through differentiation, would reflect the persistence of γH2AFX at a level below of that required to activate the recombination-dependent checkpoint.

In somatic cells, MDC1-mediated phosphorylation of H2AFX amplifies ATM response to damage. The latter promotes either the arrest of the cell cycle, or apoptosis (Stewart et al., 2003). ATM effectors on these functions include P53, which have been recently implicated in the activation of the recombination-dependent checkpoint in male meiosis (Pacheco et al., 2015, Marcet-Ortega et al., 2017). We observed that impaired co-ordination of meiotic progression and DSBs repair in *H2afx^-/-^and Mdc1^-/-^*cells at pachynema correlates with an obvious reduction of pP53 in the nucleus. Therefore, according to what is known in somatic cells, we suggest that in mouse meiosis, arrest of differentiation by the recombination-surveillance mechanism depends on the persistence, at pachynema, of a (MDC1-mediated) sufficiently high γH2AFX signal. The latter, by amplifying the ATM responses, would promote the activation of P53, thus delaying or arresting the progression as long as a sufficient number of DSBs has been repaired, and the drop of γH2AFX below a threshold level (Fig. 9).

One additional aspect of mouse meiotic cell response to DSB formation by SPO11 is the inhibition, by the ATM and MRE11 complex, of SPO11 activity (Lange et al., 2011, Pacheco et al., 2015). In somatic cells, the interaction of MRN complex with DSB promotes the recruitment of ATM onto damaged DNA before the appearance of γH2AFX. Further accumulation of the ATM and MRE11 complex around DSB sites occurs by a mechanism that requires (MDC1-mediated) H2AFX phosphorylation. Therefore, ATM and MRE11 complex-mediated control over the number of DSBs might occur either before or after phosphorylation of H2AFX. Our finding that the number of DSBs is unaltered in both *H2afx^-/-^* and *Mdc1^-/-^* cells, suggests that that γH2AFX is dispensable to exert a control over SPO11 activity (Fig. 9). Hence, while both the control of the number of DSB and the activation of recombination-dependent checkpoint depends on ATM function, the latter mechanism has distinct genetic requirements, as it is likely to need the spreading of ATM signaling within the cell, through the functions of H2AFX and MDC1.

### *H2afx* plays a minor function in the arrest of progression of zygonema cells with irreparable DSBs

In mice whose *Dmc1* has been deleted, chromosome synapsis and recombination is impaired, and no or few cells progress and incorporate H1t. Conversely, in cells where DSB formation is prevented (i.e. *Spo11^-/-^* mutant), despite defective synapsis, H1t is incorporated at high levels (Barchi et al., 2005). This indicates that the mechanism that prevents the cytological progression of zygotene stage cells to mid-pachynema, depends on the persistence of unrepaired DSBs. The genes responsible for the activation of such recombination-dependent arrest are unknown. Our observation that in a *Dmc1^-/-^* background, the lack of *H2afx* partially releases the arrest of progression of cells with persistent damage, point to a role for *H2afx* in this mechanism. However, the contribution of *H2afx* is mild, as in *Dmc1^-/-^H2afx^-/-^* mutants, cells were only faintly H1t-positive, and we only observed a modest increase in the number of H1t-positive cells, relatively to *Dmc1^-/-^*. This is likely to happen because of the persistence of a massive load of irreparable DSBs. In *Caenorhabditis Elegans*, the damage-dependent apoptosis in the germ line requires *msh4* and *msh5*, and cell death is prevented by deleting either one, regardless of the activation of the DNA-damage response (Silva et al., 2013). In mouse, albeit *Msh5* deletion prevents DSB repair in spermatocytes, the nuclei incorporate H1t at hight levels, prior to their apoptotic elimination at stage IV (Barchi et al., 2005). This makes *Msh5* an ideal candidate gene, involved in the mechanism that copes DSBs repair at zygonema with progression of spermatocytes through differentiation. While this hypothesis is tested, most likely, *Msh5* is not the only gene involved in this process. Therefore, the full understanding of the mechanisms that constrain differentiation of meiotic cells with damaged DNA continues to be a challenge for the future.

## Material and methods

### Mice and genotyping

*H2afx, Mdc1, Dmc1, p53 and Atm* mice were on a C57BL6 X 129 S/v mixed background. To minimize variability from strain background, experimental animals were compared to controls from the same litter or from the same mating involving closely related parents. Each analysis was done with 2 to 4 animals per genotype. No significant variations were observed between individual or between litter comparisons of animals with the same genotype. *H2afx^-/-^* and *Mdc1^-/-^* mice were generated from *H2afx^+/-^* and *Mdc1^+/-^* intercrosses respectively. *H2afx^-/-^Dmc1^-/-^* males were generated by corssing *H2afx^+/-^* with *Dmc1^+/-^*mice. The genotyping of *H2afx, Mdc1, Dmc1 and Atm* was performed by PCR of tail tip DNA. Primer sequences are reported elswere (Celeste et al., 2002, Lou et al., 2006, Pittman et al., 1998, Barchi et al., 2008). Tissue samples from *p53^-/-^* mice were provided by Dr. Travis Stracker (IRB).

### Preparation of meiotic chromosome spreads

Spreads of germ cell chromosomes were performed according to (Faieta et al., 2015). In brief, testes were removed from euthanized animal, decapsulated, chopped in MEM-HG, and mixed. Suspension was left to settle down and the supernatant was spun down at 7,200 rpm for 1 min. The pellet was resuspended in 0.5 M sucrose. The suspension was added to slides coated with PFA 1%, 0.015 % Triton X-100, and incubated for 2 h in a humidified chamber at room temperature (RT). At the end of incubation, slides were rinsed twice in 1:250 Photo-flo Kodak professional (no. 1464510) in water and allowed to air dry. Surface chromosome spreads were either immediately processed for immunofluorescence or stored at –80 °C for up to 6 months.

### Immunofluorescence assay and quantification of fluorescence intensity

After 10 min washes with washing Buffer 1 (0.4% Photo-flo,0.01% Triton X-100 in water), surface spreads were incubated overnight at room temperature with the primary antibody diluted in *antibody dilution buffer* (ADB) (10% goat serum, 3% bovine serum albumin [BSA], 0.05% Triton X-100 in phosphate-buffered saline [PBS]). After one wash in washing buffer 1, and one in washing buffer 2 (0.4% Kodak Photo-flo in water) for 10 min each, slides were incubated with the secondary antibody for 1 h in a pre-warmed humidified chamber at 37°C in the dark. After 10-min washes with washing buffers 1 and 2, slides were rinsed for 5 min in PBS and incubated in Hoechst/PBS solution for at least 15 minutes in humidified chamber, at RT. At the end of the incubation, slides were air-dried for 10 minutes at RT in the dark, and mounted using ProLong^®^ Gold Antifade Mountant without DAPI (Molecular Probes-Life technologies cat. num. P36934). Images were captured using Leica CTR6000 Digital Inverted Microscope connected to a charge-coupled device camera and analyzed using the Leica software LAS-AF for fluorescent microscopy.

Sources and dilutions of antibodies for immunofluorescence were as follows: mouse anti-SYCP3 (Santacruz, sc-74569) 1:300; rabbit anti-SYCP3 (Novus Biologicals nb300-231) 1:300; rabbit anti-SYCP1 (Abcam) 1:200; rabbit anti-DMC1 (Santacruz, sc 22768) 1:100; rabbit anti-γH2AX (Cell signaling) 1:400-500; guinea pig anti-H1T (a gift from M.A. Handel) 1:500; rabbit anti-pHORMAD1 s375 (a gift from C. Hoog) 1:200; rabbit anti-HORMAD2 (a gift from A.Toth) and rabbit anti-MLH3 (a gift from P. Cohen) 1:100; rabbit anti-MSH4 (Abcam ab58666) 1:75; rabbit anti-pP53 ser15 (Cell Signaling) 1:100; Monoclonal mouse anti-human Acrosin (AMC-ACRO-C5-F10-AS) 1:200; rabbit anti-BRCA1 (a gift from S. Namekawa) 1:200. Secondary antibodies from Alexa-Fluor (Molecular Probes, Life Technologies) were diluted 1:200.

The intensity of H1t and pP53 signals were measured using the LAS-AF software (Laica). Measures were performed by choosing a fix-size region of interest which included most of the diameter of the nucleus. Measures were performed only on nuclei with aproxymately the same size. For H1t measures, we considered positive, all cells with a signal above the background signal measured in pre-leptonema/leptonema nuclei.

### Terminal deoxynucleotidyl transferase dUTP nick end labeling (TUNEL)

TUNEL assay was performed folloing IF, according to (Pacheco et al., 2015), using the In Situ Cell Death Detection Kit (POD) from Roche (Cat. No. 11684817910).

### Preparation of clone DNA by Alkaline Lysis and Nick Translation

BAC clones for Y chromosome (RP24 502P5), X chromosome (RP25 470D15) and PAR region (RP24500I4) were grown in standard Luria Bertani medium at 32°C for a minimum of 16 hours. Purification of DNA, after overnight culture, was obtained with Plasmid Maxi Prep Kit (QIAGEN). The determination of DNA concentration was made by both UV spectrophotometry at 260 nm and quantitative analysis on agarose gel. Approximately 1μg of extracted BAC DNA was marked with fluorescent isotopes using Nick Translation Kit (Abbott Molecular) according to the manufacturer’s instruction. Probe sizes were determined with a quantitative analysis of agarose gel. All probes were combined with In Situ Hybridization buffer (Enzo Life Sciences) according to the manufacturer’s instruction in the presence of 0.1 μg/μl mouse Cot-1 DNA (Invitrogen). Probes were denatured at 73°C for 5 minutes in water bath.

### Fluorescence in situ hybridization (FISH) of the PAR

Following immunofluorescence, slides with nuclear spread preparation were washed with fresh PBS at RT for 5 minutes, rinsed briefly in dH_2_O, then dehydrated passing through ethanol series, and air-dried. After aging (65°C for 1 hour), slides were denatured for 7 minutes in 70%formamide/2xSSC solution at 72°C and immediately dehydrated, passing it through ethanol series at −20°C, and air-dried. The FISH probe was applied to the slides and allowed to denature in a humid chamber at 75°C for 10 minutes. Hybridization was performed in a humid chamber at 37°C for a minimum of 16 hours. Following two washes with Stringent Wash Buffer (4XSSC/0.2% Tween-20) at 55°C, slides were dehydrated through ethanol series and air-dried. Nuclei were stained with Hoechest for 20 minutes at RT. Aliquots of antifade solution were applied on the slides. Images were captured using Leica CTR6000 Digital Inverted Microscope.

### RNA extraction and RT-PCR

Testes from 10.5 and 12.5 dpp old mice for *H2afx^-/-^* and *H2afx^+/+^* were dissected and homogenized in 500 *μ* l of QIAzol Lysis Reagent (cat. no. 79306). RNA extraction was performed according to the manufacturer’s instruction and diluted in DEPC water. Contaminating genome DNA was removed with a DNA-free kit (Ambion No. AM1906). Synthesis of cDNA was performed with Invitrogen RT-PCR SuperScript III (Invitrogen, No. 18080-051) according to the manufacturer’s instruction. SPO11 splice isoforms were amplified by PCR using 2μl of cDNA. Primer sequences are listed in the supplementary material section.

### Tissue collection and processing for histological and immunofluorescence analyses

#### Histological analyses of the testis

For histological analysis, testes were collected and fixed overnight at 4°C in Bouin’s fixative (Sigma no. HT10132). Freshly collected samples were weighted before fixation. Fixed samples were embedded in paraffin. Sections of 5 μm were stained with periodic acid-Schiff/hematoxylin (PAS) (Schiff’s fuchsin-sulfite reagent, Sigma S5133). Images were captured using a Zeiss Axioskop bright field microscope equipped with a color CCD camera.

#### Immunofluorescence analyses of the testis

Testes were removed from euthanized *H2afx^+/+^, H2afx^-/-^, Mdc1^+/+^, Mdc1^-/-^, P53^+/+^, P53^-/-^* and *Atm^-/-^* adult animals and put in at least 10 volumes of 4% paraformaldehyde (PFA) in PBS, O/N at 4°C. The following day tissues were immediately processed for inclusion in paraffin blocks, as standard protocol. For Immunofluorescence analyses, paraffin embedded testes were sectioned at 3-5 *μ*m. Following deparaffinization and rehydration, sections were treated to unmask the antigenic epitope, using TrisEDTA-citrate buffer pH 7.8 (UCS diagnostic, cat. TECH199). Sections were then cooled in dH2O to continue with immunofluorescence (see above).

### Preparation of testis nuclear extracts

The procedures for the preparation of the insoluble fraction of testis nuclear extracts, and antibodies sources for the detection of the phosphorylation status of HORMAD1, HORMAD2, SYCP2, STAG3, REC8, SMC3, in *H2afx^+/+^* and *H2afx^-/-^*, are described elsewere (Fukuda et al., 2012).

### Immunoprecipitation and western blot

Immunoprecipitation and detection of pS1083 of SMC3 were performed as described in (Fukuda et al., 2012). Immunoprecipitation of SPO11 was performed using a previously described technique (Neale et al., 2005, Lange et al., 2011). In brief, testes from 12.5 dpp *H2afx^-/-^, H2afx^+/+^, Dmc1^+/+^, Dmc1^-/-^ Spo11^+/+^* and *Spo11^-/-^* mice were decapsulated and lysed in lysis buffer (1% Triton X-100, 400 mM NaCl, 25 mM HEPES-NaOH at pH 7.4, 5 mM EDTA). Lysates were subjected to two rounds of centrifugation at 13,200 rpm for 15 min each at 4° C in a benchtop centrifuge. Supernatants were incubated with monoclonal mouse anti-SPO11 antibody 180 (3 μg per pair of testes) at 4° C for 1 h, followed by the addition of 40 μI; Protein-A Agarose beads (Roche) and incubated for an additional 3 h at 4 °C. At the end of incubation, beads were washed three times with IP buffer (1 % Triton X-100, 150 mM NaCl, 15 mM Tris-HCl at pH 8.0,1 mM EDTA). Immunoprecipitates were eluted with Laemmli sample buffer. Samples were fractionated on 8 % SDS-PAGE and transferred to a PVDF membrane by the semi-dry transfer system (Bio-Rad). For western analysis, membranes were probed with antibody anti-mSPO11 antibody 180 (1:2,000 in PBS containing 0.1 % Tween 20 and 5 % non-fat dry milk) overnight at 4 °C and then with horseradish peroxidase-conjugated protein A (Abcam; 1:10,000 in PBS containing 0.1 % Tween 20 and 5 % non-fat dry milk) for 2 h at room temperature. Signals were detected using the ECL+reagent (GE Healthcare).

Western blotting of P53 was performed as described above with some modifications: testes samples collected from 10.5dpp mice were lysed in RIPA lysis buffer (100mM NaCl, 10mM MgCl_2_, 30mM Tris-Hcl, 0,5% TX-100) with the addition of 1mM DTT, 0.5mM NaOv and protease inhibitors (Roche). The anti-P53 antibody (DB-Biotech n. 002) was diluted 1:2000 in TBS-T 5% milk (ChemCruz).

### Detection of SPO11-oligonucleotide complexes

SPO11-oligonucleotide complexes were analyzed as described in (Neale et al., 2005, Lange et al., 2011). In short, following two rounds of immunoprecipitation with an anti-SPO11 monoclonal antibody (Spo11-180), SPO11-oligonucleotide complexes were labeled with [α-32P] dCTP, using terminal deoxynucleotidyl transferase (TdT), and subsequently released from the beads, by boiling it in Laemmli buffer. Sample were fractionated by SDS-PAGE, and the electrophoresed products were transferred onto a PVDF membrane. Radio-labeled species were detected using Fuji phosphor screens and analyzed with ImageGauge software. To identify SPO11 protein, the same PVDF membrane was subjected to western analysis, using the SPO11 monoclonal antibody and Protein A-HRP secondary antibody, as described above.

### Statistical analysis

Statistical analysis was performed using GraphPad Prism 5. Data were expressed as mean ± SD. The statistic significance of data expressed in percentages was analyzed using Student’s t-test previous arcsen transformation of the data (P<0.05). Statistical significance of all other data regarding the comparison of foci numbers among different genotypes was analyzed by using the Mann-Whitney test (P<0.05).

### Ethics statement

All applicable international, national, and/or Institutional guidelines for care and use of animals were followed. Mice work was approved by the animal health committee of the University of Rome Tor Vergata, and by Ministry of Health of Italy. This article does not contain any studies with human participants performed by any of the authors.

## Acknowledgments

We thank P. Cohen (Cornell University), M. A. Handel (The Jackson Laboratory), C. Höög (Karolinska Institutet), A. Toth (TU Dresden) and S. Namekawa (University of Cincinnati) for the generous gift of antibodies. We are also grateful to A. Nussenzweig (NIH) for providing *H2afx^+/-^* mice; J.C. Schimenti (Cornell University) for the *Dmc1^+/-^* mice; Sott Keeney and Julian Lange (MSKCC) for their help with the SPO11-oligo assay, and Travis Stracker (IRB) for providing samples from *p53^-/-^* testes.

## Author Summary

Meiosis is a process in which, after doubling their DNA content, germ cells undergo two rounds of division with the production of haploid gametes. As germ cells enter the first meiotic prophase, programmed double-stranded breaks (DSBs) form throughout the genome. This is a key event that – thanks to the repair of damage by homologous recombination – leads to proper synapsis of the homologs and crossover formation, with consequent segregation of the chromosomes at the end of the first meiotic division. In mammals, the number of DSBs that are produced is massive. Consequently, reduced proficiency or untimely repair of the damage can have deleterious effects on the genomic integrity of the offspring, and result in impaired production of gametes. For these reasons, germ cells are expected to have developed mechanisms that tie meiotic recombination to meiotic progression, enforce timely resolution of DSBs, and arrest differentiation when damage cannot be properly repaired. In this paper, we report that *H2afx* is needed to enforce proper and timely meiotic recombination/synapsis of the autosomes and X-Y chromosomes, and along with *Mdc1*, crossover formation. In addition, we provide compelling evidence that H2AFX and MDC1are part of the surveillance mechanism that delays or arrests the differentiation of cells with unrepaired DSBs at pachynema. Such mechanisms are expected to prevent genomic instability, with the generation of healthy offspring.

